# Hat1 Orchestrates Heterochromatin Inheritance by Regulating Localization of H3K9 Methyltransferases

**DOI:** 10.64898/2026.03.20.713225

**Authors:** Caden J. Martin, Liudmila V. Popova, Prabakaran Nagarajan, Elizabeth A. Oser, Callie M. Lovejoy, Benjamin D. Sunkel, Benjamin Z. Stanton, Michael A. Freitas, Mark R. Parthun

## Abstract

Many regions of heterochromatin associate with the nuclear periphery and are known as Lamin-associated domains (LADs). Histone acetyltransferase 1 (Hat1) is a highly conserved enzyme which acetylates newly synthesized histones H4 on lysines 5 and 12 prior to their deposition on chromatin. Hat1 is required to preserve chromatin accessibility within a subset of LADs called Hat1-dependent accessibility domains (HADs). Here we profile a diverse set of histone modifications in Hat1 KO and WT immortalized mouse embryonic fibroblasts (iMEFs) and find that Hat1 regulates diverse aspects of the structure of HADs and non-HAD LADs (nhLADS). In HADs, these changes include the conversion of H3K9me2 to H3K9me3. Analysis of H3K9-specific histone methyltransferases (HMTs) shows that that Suv39h1 and Suv39h2 have distinct localization patterns, where only Suv39h2 localizes to LADs. G9a only localizes to LADs in regions enriched for H3K9me2. We find that Hat1 loss results in a redistribution of these HMTs in both HADs and nh LADs. There is a decrease in the levels of G9a with a concomitant increase in Suv39h2. These results suggest Hat1 functions to restrain the formation of a more strongly heterochromatic state and highlight a role for Hat1 as an essential regulator of heterochromatin inheritance.

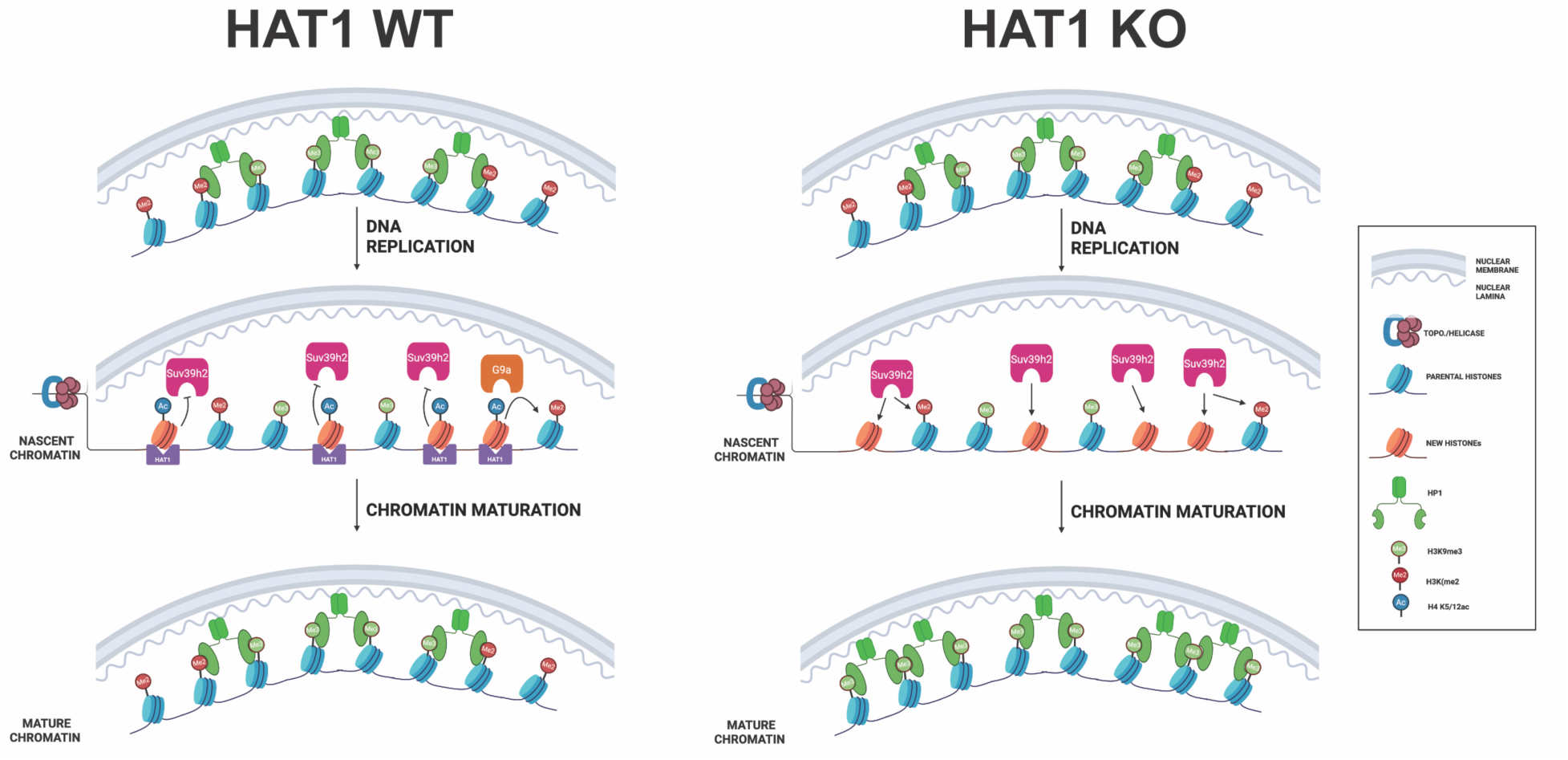

## Introduction

Eukaryotic chromatin can be broadly divided into euchromatin and heterochromatin. These two classes of chromatin perform different functions, localize differently in the nucleus, and are established and inherited by different mechanisms (1-4).

Euchromatin is gene rich, transcriptionally active, highly acetylated, and tends to localize to the interior of the nucleus, while heterochromatin is hypoacetylated, gene poor, and enforces transcriptional silencing through physical compaction and recruitment of repressive transcription factors (5,6). Heterochromatin falls into two main classes. Facultative heterochromatin is characterized by H3K27me3 and H2AK119ub and primarily silences genes in a developmental or cell cycle-specific manner(7-9). Constitutive heterochromatin is characterized by H3K9me2/3 and promotes long-term silencing of gene-poor regions, transposons, and structural elements like centromeres and telomeres(10). These histone PTMs can also recruit heterochromatin-promoting factors such as the H3K9 methylation-binding protein HP1, which promotes chromatin compaction(11,12). Both classes of heterochromatin are depleted in histone acetylation and chromatin accessibility, but these features are particularly pronounced in constitutive heterochromatin.

One way the cell maintains the structural and functional separation between euchromatin and heterochromatin is by tethering the latter to the nuclear periphery. Lamin-Associated Domains (LADs) are large chromatin domains (ranging from 10s of kilobases to megabases) which associate with the nuclear lamina by interactions with lamins and lamin-associated proteins (13). LADs contain a large fraction of the constitutive heterochromatin in the genome and are enriched in H3K9me2/3 and HP1. For example, HP1 promotes lamina association by binding histones and interacting with lamina-associated proteins like LBR and PRR14 (14,15). Other histone modifications, such as H3K9me2, H3K9me3, and H3K27me3 also influence lamina-interactions (13,16). Proper regulation of chromatin-lamina association is essential at both the cellular and organismal levels. Many LADs experience dynamic changes in lamina association through differentiation and development and vary across cell types, earning the label of facultative LADs (fLADs), in contrast to constitutive LADs (cLADs), which are always lamina-associated across cell types (17,18). Mis-regulation of lamina association can influence many features and processes, including DNA replication, genome organization, transcriptional regulation, differentiation, and cell fate commitment (19-26). Regulation of LAD chromatin is complex. Despite its repressive character, lamina association does not always correlate neatly with transcriptional silencing and LADs can contain low levels of active features like histone acetylation, often found in discrete peaks near sparse sites of accessible chromatin.

During S-phase, a cell must successfully replicate both its genome and epigenome. As the replication fork moves along the DNA, nucleosomes are disassembled from the parental strand and H3-H4 tetramers are recycled by replisome-associated chaperones onto the daughter strands near their original locations (1,27). Chromatin assembly factor 1 (CAF-1) is recruited to the replication fork by interactions with PCNA and delivers newly synthesized histones to restore nucleosome levels (28,29), resulting in roughly equal proportions of new and parental histones on the daughter strands. Many of these new histones are deposited already bearing PTMs, most notably acetylation of histone H4 lysines 5 and 12.

Following DNA replication, nucleosome organization is restored by chromatin remodelers and histone methyltransferases working to copy parental heterochromatin marks onto the new histones. Six different histone methyltransferases (HMTs) target H3K9 in mammals (30,31). Of these, Suv39h1/Suv39h2 primarily deposit H3K9me3 and G9a/GLP primarily deposit H3K9me2. Setdb1 can mono-, di-, or tri-methylate H3K9 (30,32). H3K9me1 deposited by Setdb1 helps prime nascent chromatin for further methylation by Suv39h1/2, and Setdb1 often contributes H3K9me3 at repetitive elements like endogenous retroviruses (ERVs) (33-36). Suv39h1/2 are homologs of the fission yeast HMT Clr4, which binds H3K9me3 through its chromodomain and deposits the same mark onto nearby histones in a “read-write mechanism”(37-39). This same mechanism has been demonstrated in Suv39h1 and is presumed to be shared by Suv39h2, though Suv39h1 is generally treated as the primary H3K9 trimethylase of the two (40).

Facultative heterochromatin is restored in a similar manner by cooperation between the H3K27me3 writer PRC2 and PRC1-dependent H2AK119ub (41-43). Numerous factors may influence the efficiency of the read-write mechanism, including the density of the mark in question, the 3D organization of chromatin, the presence of other modifications, linker histones, and chromatin binding proteins like HP1 (39,44-49). However, the restoration of H3K9me3 and H3K27me3 takes place following S-phase very slowly and much work remains to understand how their inheritance is regulated (27,50,51).

Histone acetyltransferase 1 (Hat1) binds newly synthesized H3-H4 dimers to acetylate histone H4 on lysines 5 and 12 before the histones are deposited onto nascent DNA by CAF-1 (52-54). Hat1 is highly conserved across eukaryotes, and its importance seems to scale with organismal complexity (53,55-57). While Hat1 deletion causes only mild phenotypes in yeast, deletion in Drosophila leads to wide-spread gene mis-regulation during development, and Hat1 KO in mice is embryonic or neonatal lethal(57,58). However, the main molecular function of Hat1 remains unclear. In addition to acetylating nascent H4, Hat1 chaperones histones and can directly associate with nascent chromatin, though it is unable to acetylate nucleosomal histones(54,59-61). Both its association with chromatin and H4K5/12ac are short-lived, disappearing from chromatin within 2 hours after replication (61-63). However, the placement of H4K5/12ac on nascent chromatin leaves them perfectly positioned to influence the early stages of chromatin maturation. Consistent with this idea, we recently demonstrated that loss of Hat1 has a significant impact on the chromatin landscape. Heterochromatin domains ranging from tens of kilobases to megabases in size displayed a loss of chromatin accessibility upon Hat1 KO, and were therefore called Hat1-dependent accessibility domains (HADs)(64). HADs are a subset of LADs and thus share their chromatin characteristics, including enrichment of H3K9 methylation, poor chromatin accessibility, and low gene density. Furthermore, nascent chromatin in Hat1 KO cells is enriched in constitutive heterochromatin associated proteins and depleted in histone acetylation relative to nascent chromatin from Hat1 WT cells, suggesting that these chromatin changes arise from loss of Hat1’s S-phase activity (61).

Here we set out to more fully understand the effects of Hat1 loss on the chromatin landscape. We find that Hat1 regulates the levels of numerous active and repressive features across LADs, and especially within HADs. HADs require Hat1 to preserve the epigenetic signature of unannotated sites which resemble primed enhancers and to maintain H3K9me2 and prevent excessive accumulation of H3K9me3 and HP1β. While other LADs do not accumulate these constitutive heterochromatin features upon Hat1 loss, they still require Hat1 to preserve both background histone acetylation and H3K27me3, a mark of facultative heterochromatin which marks LAD borders and regulates chromatin-lamina interactions. We trace the changes in H3K9 methylation to redistribution of H3K9 HMTs and show that Suv39h2, not Suv39h1, is primarily responsible for H3K9me3 in LADs. In addition, H4K5ac increases at LAD boundaries upon Hat1 loss, suggesting a mechanism by which the cell protects the surrounding genome from invasion of heterochromatin. Finally, proximity labeling demonstates that Hat1 is in proximity to nuclear lamina components, which is confirmed by proximity labeling assays with Hat1 and lamin B. These results show Hat1 to be an essential regulator of heterochromatin inheritance, most prominently in domains associated with the nuclear lamina.

## Materials and Methods

### Cell culture

Mouse embryonic fibroblasts harvested from Hat1 WT and KO mice and immortalized as previously described were cultured at 37 ⁰C in 5% CO_2_ (58). Cells were grown in Dulbecco’s Modified Eagle’s High Glucose Medium supplemented with 10% FBS, 1% glutamine, and 1% Penn-Strep and passaged approximately every 2-3 days with 0.02% Trypsin.

### Western Blot

Whole cell lysate was prepared in NP-40 Lysis Buffer and protein concentration was quantified by Nanodrop. Following electrophoresis, protein was transferred from the gel to a nitrocellulose membrane using a Bio-Rad Trans-Blot Turbo at 25 V for 20 minutes. The quality of the run and transfer was checked by visualizing the protein with Ponceau stain, which was then rinsed off with TBS-T. Membranes were blocked for 1 hr in 5% milk in TBS-T, rinsed with TBS-T, and incubated with primary antibody in 2.5% milk at 4 ⁰C overnight. Membranes were rinsed 3X for 5 min each in TBS-T, then incubated for 1 hr with HRP-conjugated secondary antibody diluted 1:5000 in 1% milk. Membranes were rinsed again 3X for 5 min in TBS-T. Pierce ECL Western Blotting Substrate (Thermo Scientific) was applied to the membrane, which was visualized using a Sapphire Biomolecular Imager (Azure Biosystems).

### CUT&Tag

CUT&Tag was performed as previously described (65) according to the EpiCypher® CUTANA™ Direct-to-PCR CUT&Tag Protocol, with a light crosslinking step adapted from (66). Briefly, 100,00 iMEFs were harvested and rinsed with PBS, then incubated in Nuclear Extraction Buffer for 10 minutes on ice. Nuclei were spun down, resuspended in PBS, and fixed by adding formaldehyde to a concentration of 0.1% and incubating 2 min at room temperature. The formaldehyde was quenched with two molar equivalents of glycine. Cells were spun and resuspended in Nuclear Extraction Buffer, then combined with 10 μL activated Concavalin A magnetic beads and incubated for 10 minutes at room temperature for nuclei to bind to the beads. The beads were separated with a magnet, resuspended in Antibody 150 Buffer with the appropriate dilution of primary antibody, and incubated overnight at 4 ⁰C. The primary antibody solution was removed and nuclei were incubated for 30 min with secondary antibody. Nuclei were rinsed twice with Digitonin 150 Buffer, then incubated 1 hr with pAG-Tn5 fusion transposase. Next, they were rinsed twice with Digitonin 300 Buffer and then incubated at 37 ⁰C for 1 hour with 10 mM MgCl_2_ to activate the Tn5 Transposase. Tagmentation was stopped by rinsing with TAPS Buffer, then samples were suspended in SDS Release Buffer and incubated at 58 ⁰C for 1 hr. SDS was quenched in 0.5% TritonX-100. DNA libraries were amplified using CUTANA High-Fidelity 2X PCR Master Mix™ with Universal i5 primer and the appropriate barcoded i7 primer, and DNA was extracted using 1.3X AMPure beads. Libraries were sequenced with paired-end Illumina sequencing.

### CUT&RUN

CUT&RUN was performed according to the EpiCypher® CUT&RUN Protocol. 500,000 iMEFs were harvested and rinsed with PBS, suspended in Nuclear Extraction Buffer (as with CUT&Tag) for 10 min on ice, rinsed with Wash Buffer, bound to activated Concavalin A beads, and incubated with primary antibody overnight at 4 ⁰C (Suv39h1 - Abcam ab283262; Suv39h2 - Abcam ab190870; G9a - Cell Signaling 68851; Setdb1 - Proteintech 11231-1-AP, Lamin B1 - Abcam ab16048). For Lamin B1 and IgG, nuclei were not extracted. 500,000 iMEFs were harvested, rinsed twice with wash buffer, bound to activated Concavalin A beads, and then incubated overnight with primary antibody at the appropriate dilution in Antibody Buffer. Samples were rinsed twice with Digitonin Buffer and incubated for 10 min with pAG-MNase. There were then rinsed twice more with Digitonin Buffer and incubated in 2 mM CaCl_2_ for 2 hrs at 4 ⁰C. Samples were quenched with 33 μL STOP Buffer containing E. coli spike-in DNA and incubated 10 min at 37 ⁰C to release fragmented DNA, then beads were separated out of solution on a magnetic rack and the supernatant was transferred to new tubes. Libraries were prepared for next generation sequencing using the CUTANA CUT&RUN Library Prep Kit.

### ChIP-seq and RNA-seq

ChIP-seq was performed as previously described (64). RNA-seq was performed in biological triplicate as previously described (67).

### Processing Sequencing Data

Raw sequencing data was analyzed to the mm10 genome with parameters *--end-to-end --very- sensitive --no-unal --no-mixed --no-discordant --phred33 -I 10 -X 700* and filtered to remove mitochondrial reads and those mapping to unlocalized or unplaced contigs. All Hat1 WT and KO libraries belonging to the same antibody/target were normalized to one another by library size. The E. coli spike-in which had been added to some CUT&RUN libraries was ignored as it did not yield consistent scaling between replicates. Scaling factors for each sample were calculated by dividing the number of filtered, mapped reads in the sample with the fewest mapped reads by the number of mapped reads in each sample. Scaling factors reads for each sample were calculated by dividing the number of filtered, mapped reads in the sample with the fewest mapped reads by the number of mapped reads in each sample. Library sizes were adjusted by subsampling with samtools view -s. Biological replicates were merged for downstream analysis and viewing using samtools merge. The normalized libraries were then converted to bedGraph and or bigwig format for viewing on IGV and other downstream analyses.

For histone PTMs, tracks showing the Hat1 KO/WT log2-fold change across the genome were created by combining the total signal for that PTM into bins across the genome using bedtools. The size of the bins varied depending on the mark and ranged from 10 kb for more diffuse or abundant marks to 300 kb for sparse marks. The WT and KO data was then loaded in R. The log2 KO/WT values were calculated for each bin, the data was filtered to remove bins with undefined log2FC values and very low signal, and a background level of reads was imputed to remaining empty bins with undefined logFC. The log2FC values were then recalculated across the genome. For some marks, a 70 kb sliding window average was used to smooth out the track and make domains clearer. For logFC tracks for CUT&RUN datasets of HMTs were created more simply using deepTools bigwigCompare with 30 kb bins.

### chromHMM

To run chromHMM on the WT and KO data together, a chromosome sizes file with “WT_” appended to the beginning of each chromosome name was combined with an identical file that had “KO_” added to each chromosome name. We went through each bam file and added “WT_” or “KO_” to the beginning of all chromosome names in each WT and KO file, respectively.

Paired WT and KO bams were then combined using samtools merge so that they could be treated as an individual sample but would map to separate chromosomes due to the modified chromosome names and could be re-separated later. Binarized files were created with 10 kb bins and the chromHMM algorithm was run for a range of states numbers. We chose to proceed with 10 states. After running the algorithm, the dense bed file was divided into two files based on WT_ and KO_ chromosome prefixes and the prefixes were removed from each file. These were used for downstream analysis in R and visualization with IGV. For analysis in R, a track with all gaps of unknown nucleotides downloaded from UCSC table browser was used to remove all bins had 50% or greater overlap with unmappable regions. All plots were created with ggplot2. Sankey plots were created using the ggsankey package.

### Data Analyses

All LAD analysis was performed using our previously published set of LADs mapped in the same cells used for this study (65). Sex chromosomes were excluded from all analyses.

For barplots of H3K4me1 (Supplemental Code S2.1) and H3K27ac (as in Supplemental Code S2.1) BED files were loaded into R and the number of peaks which overlapped the regions of interest (i.e. HADs or nhLADs) were summed for each of the 6 biological replicates (3 WT and 3 Hat1 KO). The averages across replicates for each group were plotted in ggplot2. Student’s t-test was used to calculate the p-values.

To make LAD heatmaps and profile plots, a matrix of all marks scaled across LADs was generated using deepTools computeMatrix and loaded in R (Supplemental Code S2.2). Heatmaps were made with the pheatmaps package and profile plots were made with ggplot2, except the profiles/heatmaps in Figure S1, which were made with deepTools plotHeatmap.

To calculate the fraction of lamina-associated chromatin changed for each feature, we combined the signal into 10 kb bins across the genome, calculated the log_2_ KO/WT fold-change for each bin, and selected all bins which had at least 5 kb (50%) overlap with a LAD. Bins with log_2_FC > log_2_(1.25) were considered to have gained the feature and those with log_2_FC < -log_2_(1.25) were considered to have lost it.

To determine the net change of a given feature in LADs, we calculated the total signal for each LAD using bedtools map and calculated the log_2_ KO/WT fold-change for each LAD. LADs with log_2_FC > log_2_(1.25) were considered to have gained the feature and LADs with log_2_FC < - log_2_(1.25) were considered to have lost it.

Regression analysis was performed in R and plotted using ggplot2. The r value is the Peason correlation coefficient and the p-value is the significance of the slope. Regression was performed in 3 different ways. (1) Abundance of feature 1 in 10 kb bins versus abundance of feature 2 in 10 kb bins. Regression was performed using all bins within LADs, all bins within HADs, and all bins in non-HAD LADs. Outliers were removed using a more stringent version of Tukey’s rule: bins were dropped in which the log-transformed value fell more than 4*IQR above the 3^rd^ quartile or more than 4*IQR below the 1^st^ quartile. Due to the larger number of points, the plots depict the density of points across a 2-dimensional grid. (2) Total abundance of feature 1 per LAD versus total abundance of feature 2 per LAD. Outliers were not dropped. Each point in the plot represents a LAD. (3) log_2_ KO/WT fold-change of feature 1 in LADs versus log_2_ KO/WT fold-change of feature 2 in LADs. Outliers were not dropped. Each point in the plot represents a LAD.

### APEX2 Transfections

HEK293T cells were cultured in in DMEM media (Sigma) supplemented with 10% FBS (Sigma) and 1X Penicillin/Streptomycin antibiotics (Gibco). Constructs containing pcDNA-dest40 vector and APEX2-HAT1 fusion or APEX2 only were made using the Gateway Multisite Cloning kit (Thermo Fisher). Cells were transfected with the constructs at ∼80% confluency using Lipofectamine 2000 (Life Technologies) as per manufacturer’s protocol.

### APEX2 Labeling

24 hours post-transfection cells were labeled as described with minor modifications [82]. Briefly, cells were incubated in 500uM biotin-phenol for 30 min at 37°C. Next, the cells were labeled for 1 min by addition of H2O2 to the final concentration of 1mM. Labeling reaction was quenched by freshly prepared quencher solution (10 mM sodium ascorbate, 5 mM Trolox, and 10 mM sodium azide in PBS). Cells were collected and lysed in RIPA buffer (50mM Tris-HCl, pH 7.5, 150mM NaCl, 1% NP-40, 1mM EDTA, 1mM EGTA, 0.1% SDS, and 0.5% sodium deoxycholate supplemented with 1X Complete Protease inhibitor cocktail (Roche), 1mM PMSF, 10 mM sodium azide, 10 mM sodium ascorbate, and 5 mM Trolox). To help clarify lysates, cells were sonicated with Diagenode Bioruptor Sonicator on high setting for 10 min with 30sec on/off. Additionally, cells were treated with Benzonase nuclease (Sigma) and incubated for 1 hour at 4°C with rotation and consequently centrifuged for 20 min at 20000g.

### IP and Mass Spectrometry

APEX2-MS experiment was performed in triplicate. HEK293T cells were transfected with APEX2-HAT1 fusion construct, APEX2-only construct, and untransfected cells were used as a control. APEX2-HAT1-transfected cells, APEX2-only transfected cells, and untransfected cells were treated with either biotin-phenol+H2O2 or biotin-phenol alone. Cells were labeled, and cell lysates were prepared as described above. Biotinylated proteins were isolated with streptavidin magnetic beads (Thermo Scientific). Samples were incubated with beads overnight with rotation at 4°C. After rotation, beads were captured and washed once with RIPA buffer, once with 1M KCl, once with 0.1M sodium carbonate, once with 2M urea in 10mM Tris-HCl, pH 8.0, and twice with RIPA lysis buffer. Samples were digested with trypsin via on-bead digestion overnight at 37°C and and supernatant dried by vacuum concentrator. Liquid chromatography-nanospray tandem mass spectrometry (Nano-LC/MS/MS) was performed on an Orbitrap Fusion mass spectrometer (Thermo Scientific) operated in positive ion mode. Peptides (1 µg) were separated on an easy spray nano column (PepmapTM RSLC, C18 3µ 100A, 75µm X150mm, Thermo Scientific) using a 2D RSLC HPLC system (Thermo Scientific) at 55°C. Each sample was injected into the µ-Precolumn Cartridge (Thermo Scientific) and desalted with mobile phase A (0.1% formic acid in water) for 5 minutes. Flow rate was set at 300nL/min and peptides eluted with increasing mobile phase B (0.1% formic acid in acetonitrile) over 109 min as follows: 2% to 20% in 60 min, 20-32% in 15 min, from 32-50% in 10 min, 50-95% in 5 min (holding at 95% for 2 min) and back to 2% in 2 min. The column was equilibrated in 2% of mobile phase B for 15 min before the next sample injection.

### APEX2 Data Analysis

Mass spectra from all RAW data files were converted to mzML with ProteoWizard and OpenMS (v 2.5.0(68,69). Converted files were searched on the OpenMS platform with MSGF+ search engine against a reviewed UniProt human proteome (downloaded 09/24/2020) containing the cRAP and MaxQuant contaminant FASTAs. Search parameters included: full trypsin digest, 1 missed cleavage, oxidation of methionine and acetylation of lysine as variable modifications, precursor mass tolerance 20 ppm and fragment mass tolerance 0.8 Da. Testing for differentially expressed proteins was performed in R with the limma package version 3.42.2 using an imputation method described in Gardner and Freitas (70). Briefly, samples were selected for pair-wise comparisons prior to filtering out lowly expressed proteins, and missing values were imputed with a multiple imputation approach by treatment group. Data was quantile normalized, and significance (p-value < 0.05) determined by a modified exact test. To correct for background biotinylation, significantly biotinylated proteins were identified in APEX2-Hat1 biotin-phenol+H2O2 (BPH2O2) condition compared to APEX2-Hat1 biotin-phenol only (BP) condition. Next, significantly biotinylated proteins were identified in APEX2-only BPH2O2 condition compared to APEX2-only BP condition. Also, significantly biotinylated proteins were identified in untransfected BPH2O2 condition compared to untransfected BP condition. Finally, proteins found to be significantly biotinylated in the two latter conditions were removed from the list of proteins significantly biotinylated with APEX2-Hat1 construct.

### Proximity Ligation Assays

Three Hat1 WTor Hat1 KO MEF cell lines were seeded in equal quantities on coverslips and allowed to attach for 24 h. Cells were then permeabilized with 0.5% Triton X-100 and fixed with 4% PFA simultaneously for 15 min, rinsed with PBS, and fixed again with 4% PFA for 10 min at room temperature. After several PBS washes cells were blocked with 5% BSA for 1 h at room temperature. BSA was removed with PBS and primary antibodies detecting two proteins of interest were diluted in 1% BSA, 0.3% Triton X-100 and added to cells overnight at 4 °C (mouse anti-LaminB1, 1:200, Abcam ab8982; rabbit anti-LaminB1, 1:200, Abcam ab65986; rabbit anti-HAT1, 1:1000, Abcam ab193097). The following day, primary antibodies were removed with PBS and cells were subjected to the Duolink^TM^ Proximity Ligation Assay protocol according to the manufacturer’s instructions (Sigma: DUO92008, DUO92004, DUO92002, and DUO82049). After amplification, nuclei were stained with 20 mM Hoechst 33342 Fluorescent Stain and mounted on slides using Vectashield. Slides were analyzed under a Zeiss LSM 900 Airyscan 2 Point Scanning Confocal microscope. Images were acquired using Zen Blue 3.0 and quantification was completed using ImageJ version 1.52t according to a previously described protocol (71).

**Table 2.1.** Antibodies used in experiments.

| **Antigen** | **Company** | **Catalog Number** | **Application** |
| --- | --- | --- | --- |
| HP1β | Cell Signaling | 8676 | ChIP-seq, Western Blot |
| H3K9me3 | Abcam | ab8898 | CUT&Tag |
| H3K9me2 | Abcam | ab1220 | CUT&Tag |
| H3K27me3 | Cell Signaling | 9733 | CUT&Tag |
| H3K4me1 | Cell Signaling | 5326 | CUT&Tag |
| H3K9ac | Cell Signaling | 9649 | CUT&Tag |
| H3K14ac | Abcam | ab52946 | CUT&Tag |
| H3K27ac | Cell Signaling | 8173 | CUT&Tag |
| H4K5ac | Abcam | ab51997 | CUT&Tag |
| H4K12ac | Abcam | ab46983 | CUT&Tag |
| Lamin B1 | Abcam | ab16048 | CUT&RUN |
| Lamin B1 | Abcam | ab8982 | PLA |
| Lamin B1 | Abcam | ab65986 | PLA |
| Suv39h1 | Abcam | ab283262 | CUT&RUN, Western Blot |
| Suv39h2 | Abcam | ab190870 | CUT&RUN, Western Blot |
| G9a | Cell Signaling | 68851 | CUT&RUN, Western Blot |
| Setdb1 | Proteintech | 11231-1-AP | CUT&RUN, Western Blot |
| IgG | Epcypher | 13-0042 | CUT&Tag, CUT&RUN |
| HP1α | Cell Signaling | 2616 | Western Blot |
| HP1γ | Invitrogen | MA3-054 | Western Blot |
| B-Actin | Cell Signaling | 3700 | Western Blot |
| GAPDH | Cell Signaling | 5174 | Western Blot |
| Hat1 | Abcam | ab193097 | Western Blot, PLA |

## Results

### Hat1 Regulates Accessibility at Primed Enhancers in Heterochromatin

Previous results showed that Hat1 was required to preserve peaks of chromatin accessibility within a subset of Lamin-associated chromatin that were termed Hat1-dependent accessibility domains (HADs). Given the highly repressed and condensed nature of constitutive heterochromatin, these peaks of accessibility are sparsely distributed and their role is unclear. Comparison with publicly available datasets suggested that they overlap with histone modifications characteristic of active enhancers (H3K4me1 and H3K27ac). However, they do not coincide with previously known enhancers (64).

To test whether the heterochromatic peaks of accessibility overlap with these enhancer marks in our cell lines and whether those modifications are regulated by Hat1, we performed CUT&Tag in immortalized mouse embryonic fibroblasts (iMEFs) to detect the general enhancer mark H3K4me1 and the active enhancer mark H3K27ac (72). Just as we saw previously with chromatin accessibility, we observed a net loss of H3K4me1 peaks in Hat1 KO HADs compared to WT HADs (Figure 1A). The effect of Hat1 on H3K4me1 was specific to HADs as there was no overall loss of H3K4me1 in non-HAD LADs (nhLADs) or in inter-LADs (Figure 1B and Figure S1A,B). While some peaks of H3K27ac were also lost, we did not see a net reduction of H3K27ac peaks across HADs, nhLADs, or inter-LADs (Figure 1B and Figure S1A,C). Biological variability between replicates from different mice prevented accurate quantification of the overlap between H3K4me1 peaks and lost ATAC-seq peaks, but visual comparison showed that many of the lost peaks of accessibility did overlap with peaks of H3K4me1, while a much smaller number also overlapped with H3K27ac. Figure 1C shows a genome browser view of a 175 kb region of a HAD on chromosome 1 that contains the Serpin B8 gene. Consistent with the designation of this region as a HAD, there were distinct peaks of ATAC-seq signal that were present in Hat1 WT cells that were absent in Hat1 KO cells. The most prominent of these peaks was located just upstream of the Serpin B8 gene. Notably, this Hat1-dependent site of accessibility co-localized with a peak of H3K4me1 that was also Hat1-dependent. While H3K27ac was present upstream of Serpin B8, the signal was more dispersed and was not as obviously Hat1-dependent. Consistent with the hypothesis that Hat1 can regulate heterochromatic enhancers, RNA-seq analysis indicated that there was a significant Hat1-dependent decrease in Serpin B8 mRNA.

**Figure 1.**
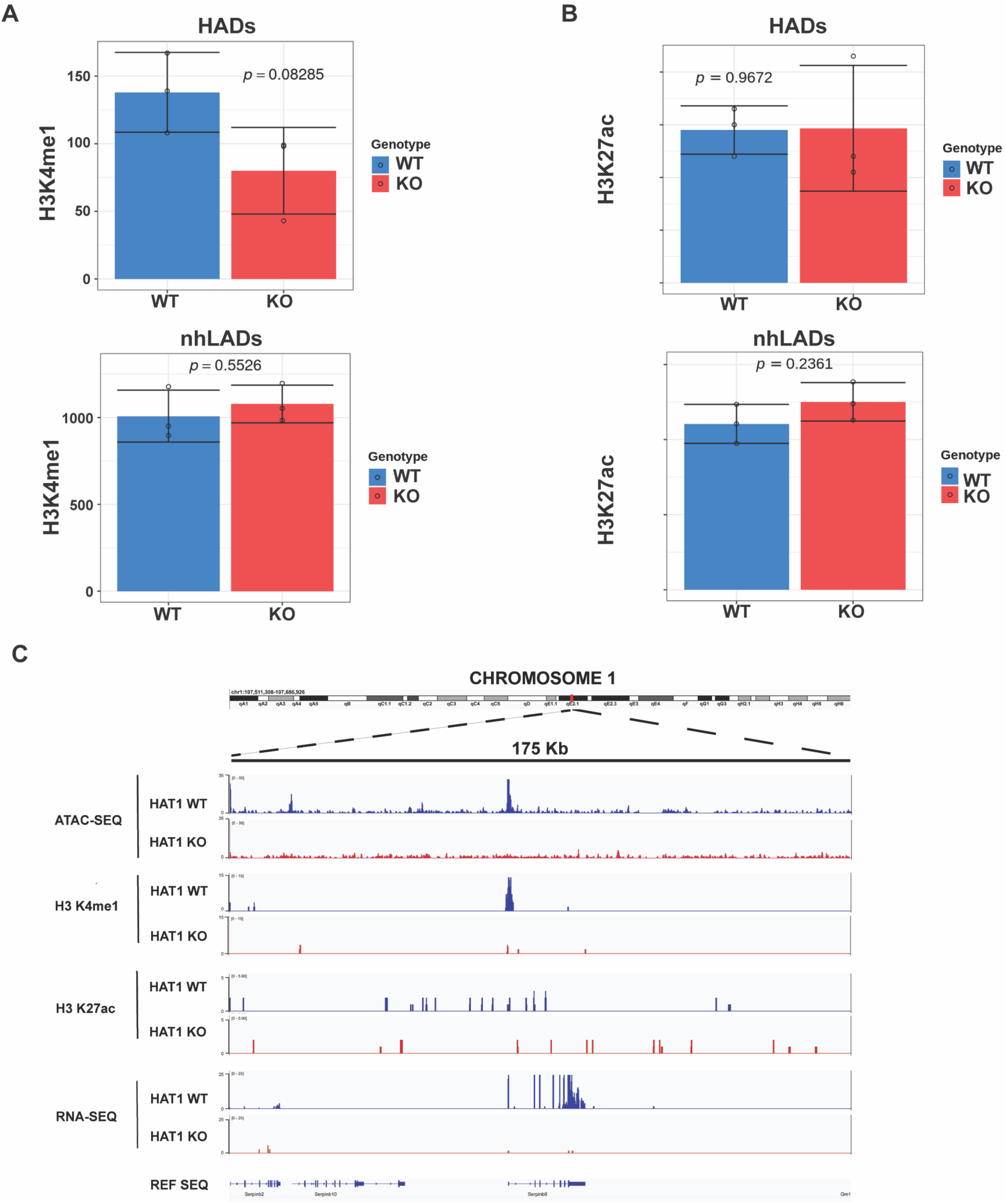
Hat1 regulates primed enhancers in HADs. (A) Average number of H3K4me1 peaks between Hat1 KO and WT cells in HADs (top) and non-HAD LADs (bottom). Error bars represent 1 standard deviation. P-values were calculated by Student’s t-test with n=3 biological replicates. (B) Comparison of H3K27ac as in panel A. (C) IGV browser view of chromatin accessibility, H3K4me1, H3K27ac, and RNA-seq at the Serpin B8 locus.

### Hat1 preserves basal acetylation across LADs

Hat1 clearly impacts sites of chromatin accessibility that possess enhancer characteristics in constitutive heterochromatin. To determine whether the effect of Hat1 is restricted to these discrete loci or whether Hat1 more broadly regulates the structure of chromatin domains associated with the nuclear lamina, we determined the genome-wide distribution of a wide variety of histone PTMs in Hat1 WT and Hat1 KO iMEFs. We performed CUT&Tag for several sites of acetylation on histones H3 and H4. These included H4K5ac and H4K12ac, which are direct targets of Hat1, as well as H3K9ac, H3K14ac, H3K27ac. Note that H4K5/12ac can also be deposited in a non-replication-dependent manner by histone acetyltransferases other than Hat1.

Calculation of the Hat1 KO/WT log_2_ fold-change showed a consistent loss of histone acetylation across not only HADs, but almost all lamina-associated domains (Figure 2A). The exception to this rule was H3K14ac. This modification showed a significant reduction in HADs and a more variable response across nhLADs.

**Figure 2.**
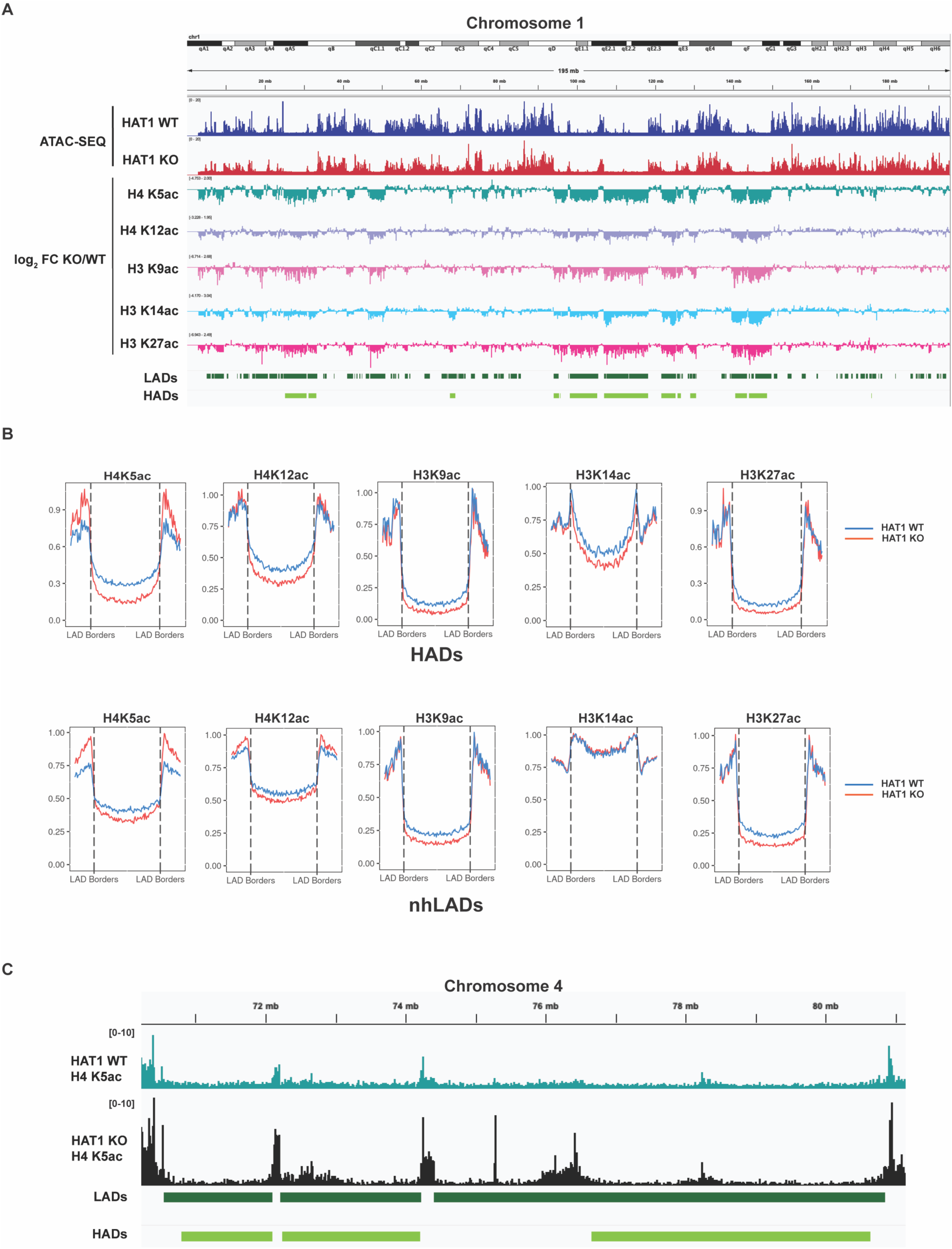
Acetylation changes in LADs upon Hat1 loss. (A) IGV browser view of chromatin accessibility and the Hat1 KO/WT log_2_ fold-change of histone acetylation across chromosome 1. (B) Profile plots depicting the abundance of histone acetylation in Hat1 WT (blue) and KO (red) cells across LADs that overlap HADs (“HADs”) and LADs which do not overlap HADs (“nhLADs”). Profiles show all LADs scaled to the same size and extend 300 kb beyond each LAD border. (C) IGV browser view of a region in chromosome 4 showing loss of basal H4K5ac across LADs and gain of H4K5ac outside LAD borders in Hat1 KO cells.

To visualize the levels of each site of histone acetylation across all HADs and nhLADs, we scaled each HAD/nhLAD to a uniform size and plotted the average signal for each acetylation site across them (plus 300 kb up- and down-stream of the HAD/nhLAD borders). For the HAD plots, we used the set of LADs which overlap HADs rather than our previously determined HAD set itself (64). This was for better visualization and clearer comparison. The boundaries of LADs can be mapped more precisely than HADs since the increased signal density in domains of lamina association are more easily measured than loss of sparse accessibility peaks. For the rest of this paper, these HAD-overlapping LADs will simply be referred to as HADs. The HAD/nhLAD profiles are shown in Figure 2B. In Hat1 WT cells, H4K5ac, H4K12ac, H3K9ac, and H3K27ac were present at uniformly low levels across the entirety of both HADs and nhLADs. The levels of these sites of acetylation rose dramatically at the border of the HADs/nhLADs and peaked just outside the HAD/nhLAD borders.

Despite this relatively low abundance in HADs/nhLADS, loss of Hat1 resulted in a significant decrease in H4K5ac, H4K12ac, H3K9ac, and H3K27ac across the length of HADs/nhLADs (Figure 2B). The H4K12ac, H3K9ac, and H3K27ac profiles changed only within LADs. The peak observed just outside the HAD/nhLAD border was indistinguishable between Hat1 WT and Hat1 KO cells. In contrast, In Hat1 KO cells there was a marked increase in the level of H4K5ac just outside of HAD/nhLAD borders Figure 2B and 2C).

As we recently reported, the pattern observed for H3K14ac was quite distinct and differentiates HADs from nhLADs. In Hat1 WT cells, there was a large peak of H3K14ac just inside the HAD borders that was much higher than the average levels of H3K14ac seen throughout euchromatin. The level of H3K14ac then gradually decreases throughout HADs to levels somewhat lower than those seen in euchromatin. In nhLADs, the peak just inside the nhLAD border is maintained but there is a much smaller decrease in the average level of H3K14ac across the length of the nhLAD, with the result that the overall level of H3K14ac across nhLADs was higher than the average levels seen in euchromatin. In HADs, loss of Hat1 resulted in a decreased level of H3K14ac across the entirety of the HAD. However, in nhLADs there was no change in H3K14ac. This supports the classification of HADs as a distinct set of lamina-associated regions of the genome that are specifically regulated by Hat1.

Very little change in histone acetylation was seen in euchromatic areas, indicating that Hat1 primarily regulates chromatin associated with the nuclear lamina. 948 LADs (84.1%), or over 95% of lamina-associated chromatin, experienced reduction of one or more acetylation marks upon Hat1 loss. H3K9ac and H3K27ac were the most depleted, being reduced in 64.0% and 72.5% of LADs, respectively, while H4K5ac and H4K12ac were lost in 54.0% and 45.8% of LADs, respectively. Looking specifically at HADs, the change was even greater, with 173 out of 180 HADs experiencing reduction of one or more acetylation mark.

The reduced histone acetylation in Hat1 KO LADs is unlikely to be caused by reduced accessibility of Hat1 KO chromatin to the pAG-Tn5 transposase, since a control with IgG in place of primary antibody shows only very minor loss of background signal in the absence of Hat1, even in LADs. HADs did show a slight reduction in IgG signal in Hat1 KO cells, but it is not nearly sufficient to explain the reduction in histone PTM signal observed (Figure S2A,B).

### Hat1 selectively regulates constitutive heterochromatin components in HADs and nhLADs

The analysis of the impact of Hat1 loss on global histone acetylation suggests that Hat1 regulates the chromatin state of most or all LADs, not only the subset identified as HADs. Informed by previous ChIP-seq data which had shown that Hat1 KO increased the density of H3K9me3 in HADs (64), we hypothesized that the loss of histone acetylation in LADs was a downstream result of increased constitutive heterochromatin features leading to a more repressive chromatin state in LADs.

To gain a more comprehensive picture of the role of Hat1 in regulating repressive chromatin structure in HADs and nhLAD, we used CUT&Tag to map H3K9me2, H3K9me3, and H3K27me3 in Hat1 WT and Hat1 KO cells. While H3K9me2 and H3K9me3 are thought to characterize constitutive heterochromatin and H3K27me3 is linked to facultative heterochromatin, all of these repressive histone methylations are associated with lamina-associated domains. In fact, LADs can be divided into 3 distinct clusters based on which is the dominant methylation in a given LAD (Martin et al, 2025).

Comparing the patterns of methylation between HADs and nhLADs identified several Hat1-dependent alterations that differ between HADs and LADs (Figure 3A). As expected, profiles of H3K9me3 show that it was highly enriched in both HADs and nhLADs. H3K9me3 formed a broad plateau spanning most of the HAD or nhLAD, then dropped sharply inside the HAD/nhLAD borders. The level of H3K9me3 in HADs increased in Hat1 KO cells while it slightly decreased in nhLADs, consistent with our previous results (64). Though also characteristically enriched in LADs, H3K9me2 displayed a very different profile. H3K9me2 enrichment peaks just inside both HAD and nhLAD borders. However, while loss of Hat1 has little net effect on H3K9me2 in nhLADs, there is a dramatic Hat1-dependent loss of H3K9me2 in HADs. As reported previously, H3K27me3 was found in a sharp peak at the border of both HADs and nhLADs. Levels of H3K27me3 were lower throughout the LAD interior in both HADs and nhLADs, especially the former. In both groups, loss of Hat1 resulted in reduced H3K27me3 levels across the domains. We saw a notable reduction of H3K27me3 in about 75% of LADs, suggesting that Hat1 plays a fundamental role in regulating H3K27me3 throughout lamina-associated chromatin. Notably, the changes in histone methylation were not strictly confined to HADs (Figure S3A). For example, many nhLADs also gained H3K9me3, but more often they lost it, resulting in a net reduction across nhLADs.

**Figure 3.**
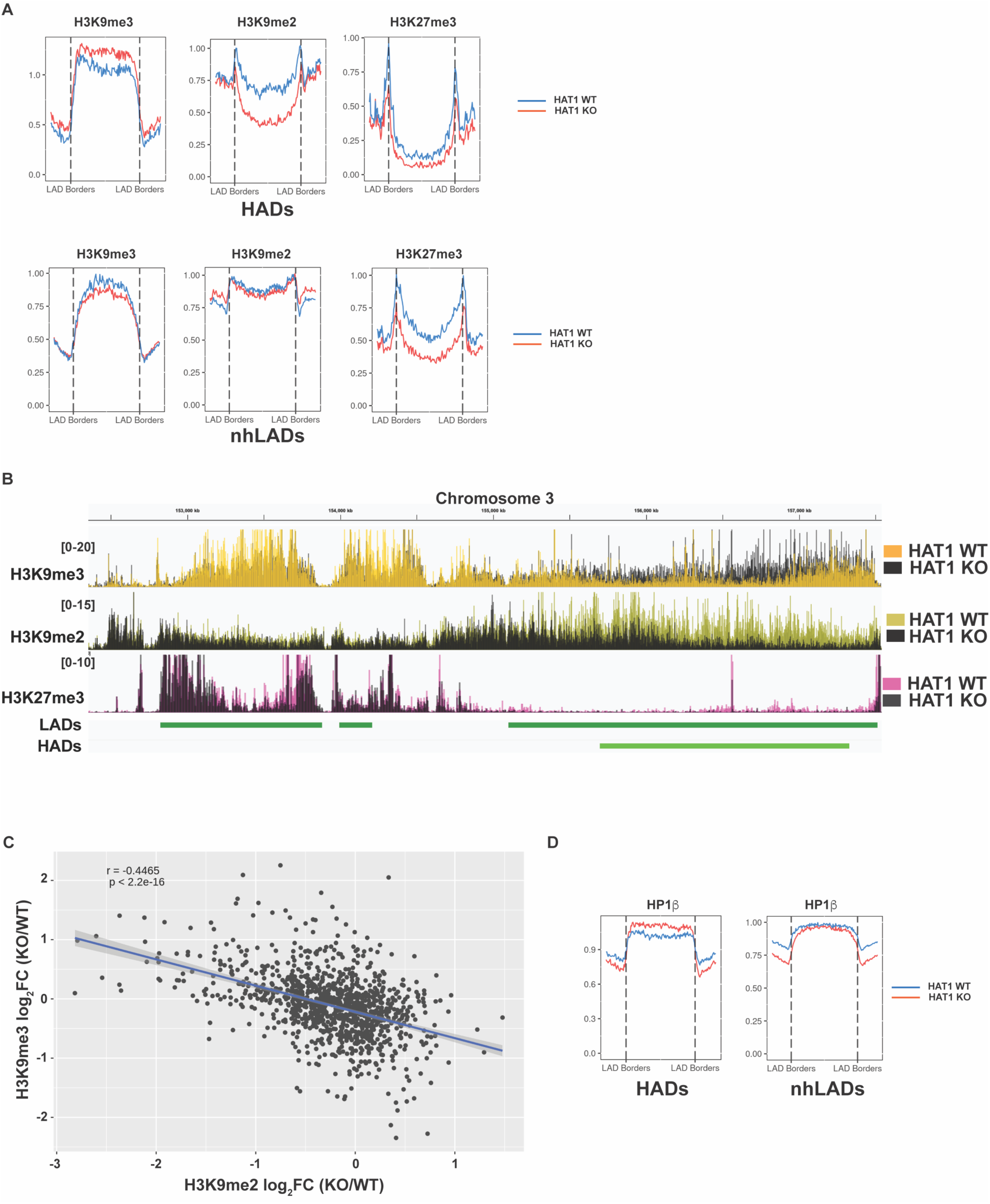
Heterochromatin changes upon Hat1 loss. (A) Profile plots depicting the abundance of the specified PTM across LADs in Hat1 WT (blue) and KO (red) cells. Profiles show all LADs scaled to the same size and extend 300 kb beyond each LAD border. (B) IGV browser view of H3K9me2, H3K9me3, and H3K27me3 CUT&Tag in 3 LADS, one of which overlaps a HAD. (C) Regression plot of H3K9me2 logFC vs. H3K9me3 logFC in LADs. Each LAD is a single data point. (D) Profile plots like those in panel A, comparing HP1β in LADs that overlap HADs and LADs which do not overlap HADs.

Viewing H3K9me3, H3K9me2, and H3K27me3 on a genome browser highlights the distinct chromatin structures in HADs and nhLADs and suggests that the opposite effects of Hat1 loss on H3K9me2 and H3K9me3 levels may be linked. Figure 3B shows a ∼5 Mb region of chromosome 3 that contains 3 LADs, one of which overlaps a HAD. In the nhLADs, no Hat1-dependent changes in H3K9me3 or H3K9me2 were observed. H3K27me3 was present at higher levels in the nhLADs than the HAD, and experienced a Hat1-dependent decrease in H3K27me3 across both. The HAD displayed a clear Hat1-dependent increase in H3K9me3 that occurs in the same region where there is a Hat1-dependent loss of H3K9me2. The opposing effects of Hat1 on H3K9me2 and H3K9me3 can be observed across all LADs, as a regression analysis of the log_2_ fold change of these marks in Hat1 WT and Hat1 KO cells shows a marked negative correlation (Figure 3C).

As the primary structural component of constitutive heterochromatin that binds to H3K9me2/3, we wanted to determine whether the association of HP1 with HADs and nhLADs is influenced by Hat1. We used ChIP-seq to measure the genome-wide distribution of HP1β. We generated profiles of the average signal of HP1β across HADs and nhLADs as done for the histone PTMs. As shown in Figure 3D, there is an increase in HP1β in HADs but a small decrease in HP1β in nhLADs, consistent with the HAD-specific increase in H3K9me3 observed in HADs. Western blot analysis found no expression changes in any HP1 isoform in Hat1 KO cells (Figure S3B-D), suggesting that HP1β is redistributed to HADs from other regions of the genome upon Hat1 KO. This may help explain the loss of HP1β in other LADs. Interestingly, the increases in H3K9me3 and HP1β in HADs seem to be independent of one another, as there is an insignificant degree of overlap between the sets of HADs that gain them and no correlation between their respective log_2_ fold-changes (Figure S3E). Together, these data suggest that Hat1 regulates the patterns of repressive histone methylation in lamina-associated chromatin, including the conversion of H3K9me2 to H3K9me3 and a concomitant increase in HP1β in HADs.

### Hat1 regulates the distribution of H3K9-specific HMTs in LADs

One mechanism by which Hat1 may influence the distribution of H3K9me3 and H3K9me2 throughout lamina-associated constitutive heterochromatin is through regulating H3K9-specific histone methyltransferases (HMTs). Hat1-dependent acetylation has been shown to control the stability of several proteins by regulating protein ubiquitylation and proteosome-mediated degradation (73). However, Western blot analysis demonstrated that loss of Hat1 did not alter the abundance of the H3K9-specific HMTs Suv39h1, Suv39h2, G9a, or Setdb1 (Figure S4A-C).

We next determined whether Hat1 regulates the genome-wide distribution of H3 K9-specific HMTs. We used CUT&RUN to profile Suv39h1, Suv39h2, G9a, and Setdb1 in the same HAT1 KO and WT iMEF cells lines that were used to profile the histone PTMs described above. In Hat1 WT cells, contrary to expectations, we found that Suv39h1 did not coincide with the broad domains of H3K9me3 found in LADs. Rather, Suv39h1 localized primarily to discrete peaks outside of LADs, often in euchromatic regions, and was largely absent from most LADs (Figure 4A and 4B). In contrast, while Suv39h2 colocalized with Suv39h1 in a subset of peaks in euchromatin, it was primarily found in broad domains that corresponded to domains of H3K9me3 in LADs (Figure 4A and 4B). Regression analysis showed that the abundance of Suv39h2 in LADs was closely correlated with the level of H3K9me3, while there is little correlation between H3K9me3 and Suv39h1 in LADs (Figure 4C). Setdb1 also tended to be depleted in LADs, resembling the distribution of Suv39h1. Regression analysis indicated that there is no correlation between H3K9me3 in LADs and the level of Setdb1 (Figure 4C).

**Figure 4.**
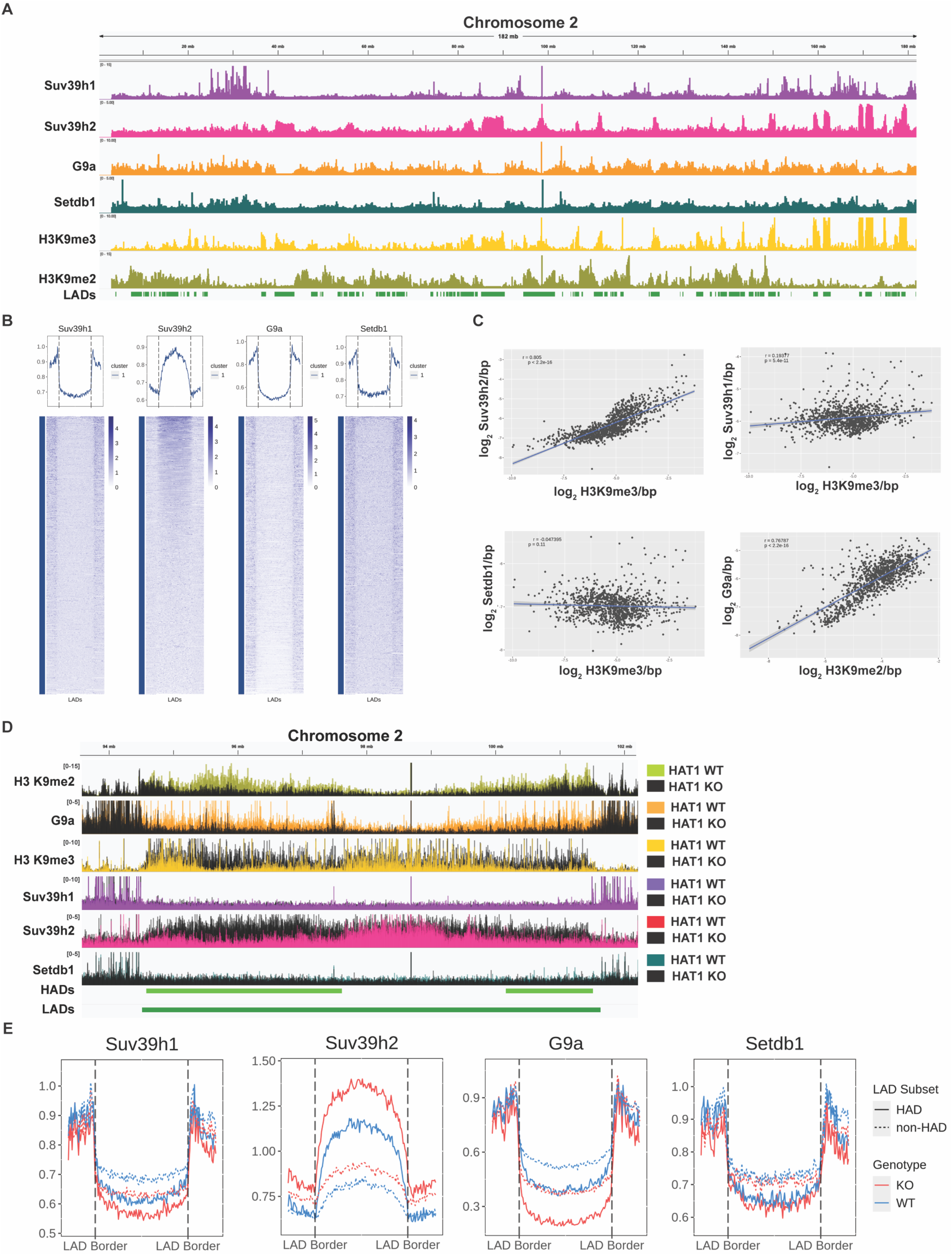
H3K9 histone methyltransferases in Hat1 KO and WT cells. (A) IGV browser view of HMTs and H3K9me2/3 across chromosome 2. (B) Profiles of HMT abundance across LADs. Profiles show all LADs scaled to the same size and extend 300 kb beyond each LAD border. (C) Regression plots comparing the log-transformed abundance of H3K9me3 vs. Suv39h2, Suv39h1, and Setdb1 in LADs and of H3K9me2 vs. G9a in LADs. Each LAD is a single data point. (D) IGV browser view illustrating changes in H3K9me2/3 and HMTs across a single LAD in Hat1 WT (colored) and KO (black). HADs overlap both ends of the LAD. (E) Profile plots of scaled LADs comparing the abundance of each HMT in Hat1 WT (blue) and KO (red) in HAD-overlapping LADs (solid lines) and non-HAD LADs (dotted lines).

G9a was generally depleted in LADs, like Suv39h1 and Setdb1. However, regions of G9a accumulation often appeared inside LADs which showed enrichment inside the border and decreased toward the LAD interior (Figure 4A). This pattern was highly similar to that observed for H3K9me2. Indeed, we found that G9a was highly correlated with H3K9me2 inside LADs, but not correlated outside LADs (Figure S4D). Interestingly, the level of Suv39h2 across individual LADs often appeared negatively correlated with that of G9a. Linear regression confirmed this negative correlation (Figure S4E), while the opposite trend was seen in inter-LAD regions. Not only was Suv39h2 generally lowest near the borders, where G9a was the highest, but peaks of G9a observed near the center of LADs often corresponded with a dip in an otherwise uniform distribution of Suv39h2 (Figure 4D). These results suggest that, in iMEFs, Suv39h2 is primarily responsible for H3K9me3 in LADs and G9a is primarily responsible for H3K9me2 in LADs.

Comparing HMT levels in Hat1 WT and KO cells, we found that Hat1 loss significantly increased Suv39h2 levels in HADs, while G9a levels were reduced. Changes in in the levels of Suv39h2 and G9a corresponded to changes in H3K9me3 and H3K9me2, respectively. This is demonstrated by the genome browser view shown in Figure 4D, which displays an ∼9 Mb section of chromosome 2. This region contains a large LAD with HADs located at each end. In the WT cells, there were high levels of H3K9me2 inside the borders of the LAD, in the regions designated as HADs. In Hat1 KO cells, the levels of both H3K9me2 and G9a drop significantly. These regions also showed corresponding increases in H3K9me3 and Suv39h2. There was little or no change in the levels of either Suv39h1 or Setdb1.

Hat1-dependent changes in in H3K9-specific HMTs were not limited to HADs, though they were most pronounced there, on average (Figure 4E, Figure S4F). We profiled the localization of Suv39h1, Suv39h2, G9a, and Setdb1 in Hat1 WT and Hat1 KO cells across all LADs. These profiles revealed changes across most LADs, as well as distinct characteristics of HADs. While the level of Suv39h1 was low across all LADs, it was higher in nhLADs relative to HADs. Loss of Hat1 reduced the level of Suv39h1 across both nhLADs and HADs, similar to what was observed for H4K5/12ac and H3K9/27ac. The level of Suv39h2 is much higher in HADs than nhLADs. Hat1 KO increases Suv39h2 levels in both, but especially HADs. Conversely, G9a levels were higher in nhLADs than HADs, and decreased in both in Hat1 KO cells. The level of Setdb1 was higher in nhLADs than HADs but Hat1 had no effect on Setdb1 localization.

The Hat1-dependent changes in HMT localization to LADs may be linked to the Hat1-dependent changes in histone methylation observed in these domains. Regression analyses show there was a strong correlation between Hat1-dependent changes in G9a and H3K9me2 (Figure S5A).

Similarly, there was a strong correlation between changes in Suv39h2 and changes in H3K9me3 (Figure S5B). There was no correlation between Hat1-dependent changes in Suv39h1 or Setdb1 and H3K9me3 (Figure S5C,D) In addition, the Hat1-dependent loss of G9a may be linked to the increase in Suv39h2. Regression analysis of the change in G9a and Suv39h2 showed a strongly negative correlation (Figure S5E). Together these results show that Hat1 influences the patterns of H3K9 methylation in LADs by controlling the localization of histone methyltransferases.

### chromHMM reveals that Hat1 restrains a shift to more strongly heterochromatic states

ChromHMM is a Hidden Markov Model-based chromatin classification tool which groups regions of the genome into states based on the adjacent states and which features are enriched. To more clearly see how Hat1 alters the character of chromatin across the genome, we ran chromHMM on our genomic maps of Lamin B1, HP1β, histone methyltransferases, and histone modifications using 10 kb bins with a 10 state model (Figure 5A). The model was trained on the Hat1 WT and Hat1 KO data together to ensure the states were the same and the two conditions could be compared.

**Figure 5.**
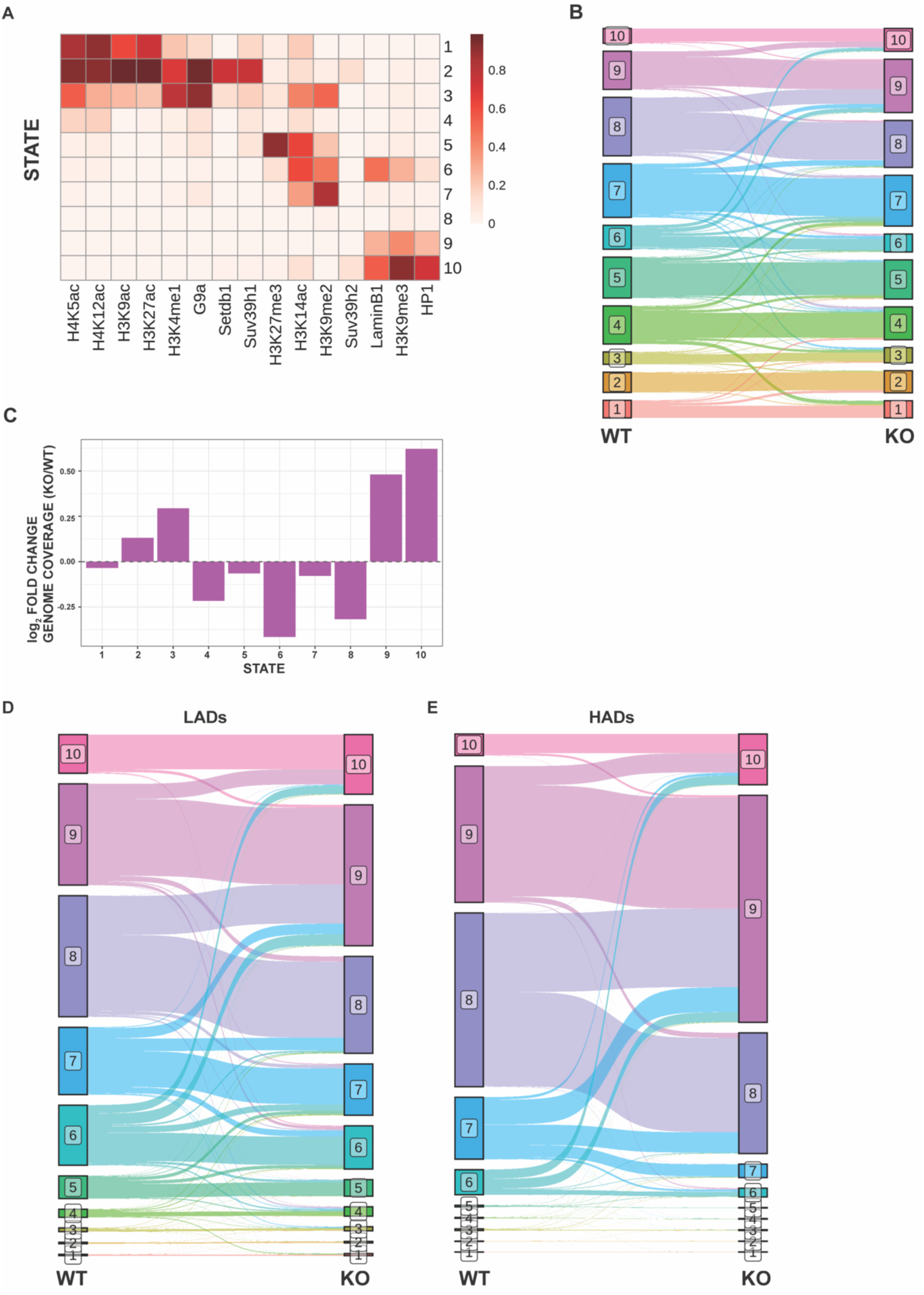
ChromHMM comparison of Hat1 KO and WT chromatin. (A) States output by chromHMM in combined WT and KO genome. The heatmap shows the probability of the mark being present in a bin of that state. (B) Sankey plot showing state changes for 10 kb bins across the genome between WT and Hat1 KO cells. (C) Change in total genome coverage for each state (log_2_ KO/WT calculated from the number of bins assigned to that state in the WT and KO genomes). (D-E) Sankey plots showing state changes for 10 kb bins in LADs and HADs.

The output states formed a rough continuum from euchromatin enriched in histone acetylation (states 1 and 2) to constitutive heterochromatin-enriched and lamina-associated (states 9 and 10). State 3 appeared to be an intermediate state that was similar to states 1 and 2 but was also enriched for H3K14ac and H3K9me2. States 5-7 were enriched to varying degrees in H3K27me3, H3K14ac, and H3K9me2, marks enriched inside LAD borders and across some LAD bodies. States 4 and 8 show very low enrichment of all features due to the nature of the algorithm used in chromHMM, which categorizes marks as simply present or absent in a given bin, placing many bins with infrequent enrichment of marks in apparently featureless states. The transitions matrix showed that state 4 usually represented euchromatic regions of low feature enrichment, while state 8 represented repressive regions with low enrichment of heterochromatin features.

Across the genome as a whole, Hat1 KO chromatin showed a significant increase in the abundance of states 9 and 10, corresponding to constitutive heterochromatin, with milder increases in euchromatic states 2 and 3. There was a notable reduction in states 6 and 8, which represent a more moderate constitutive heterochromatin signature (Figure 5B and 5C).

WT LADs were predominantly enriched in states 6 – 9. The proportion of LAD area occupied by states 6 – 10 remained similar in Hat1 KO cells, but there was a general transition towards higher states, with states 9 and 10, which are found in the interior of LADs, experiencing the greatest relative growth at the expense of states 6 – 8, which predominate near LAD borders (Figure 5D). A similar shift was seen in HADs, which underwent a significant movement from states 7 – 9 to states 9 and 10, especially 9 (Figure 5E). These changes indicate that loss of Hat1 results in a shift of existing heterochromatin states to states with greater enrichment of repressive constitutive heterochromatin features.

### Hat1 colocalizes with Nuclear Lamina Proteins

Our genomic analyses indicate that the most prominent effect of Hat1 is on nuclear lamina-associated domains of constitutive heterochromatin. To determine whether HAT1 is directly linked to the nuclear lamina, we used APEX2-based proximity labeling to identify proteins that localize near HAT1. APEX2 is an ascorbate peroxidase that generates a very short-lived biotin-phenoxyl radical in the presence of biotin phenol and H2O2 that can modify proteins in close proximity (Hung et al., 2016)(74,75). We transfected HEK293T cells with constructs that expressed either an APEX2-HAT1 fusion protein or APEX2 alone to control for non-specific biotinylation. To confirm that the APEX2-Hat1 construct exhibited a pattern of localization similar to that of endogenous HAT, we performed subcellular fractionation. Upon staining with an anti-HAT1 antibody, we observed that the APEX2-HAT1 fusion was localized to both the cytoplasm and the nucleus, similar to the localization of endogenous HAT1. Importantly, the APEX2-HAT1 fusion protein was expressed at a level comparable to endogenous HAT1 in both the nucleus and the cytoplasm (Supplementary Figure S6).

Following incubation of cells with biotin phenol and H2O2, extracts were generated, biotinylated proteins were isolated by streptavidin purification and identified by mass spectrometry. A total of 29 proteins were found to be significantly biotinylated by APEX2-HAT1 after filtering out the proteins that were significantly biotinylated in control cells (untransfected cells and cells expressing APEX2 only). A volcano plot of significantly biotinylated proteins included HAT1 and histone H4, supporting the validity of this approach (Figure 6A). PCNA was also significantly biotinylated, consistent with recent reports demonstrating that HAT1 transiently localizes to newly replicated DNA (Agudelo Garcia et al., 2017; Agudelo Garcia et al., 2020a; Nagarajan et al., 2013). Importantly, LBR (lamin B receptor) and EMD (emerin), a protein that links the nuclear lamina to the nuclear membrane, were also found to be significantly biotinylated by APEX2-HAT1. The identification of LBR in proximity to Hat1 I consistent with the reent identification of LBR as a direct substrate of Hat1(76). These results suggest that a population of HAT1 resides in proximity to the nuclear lamina.

**Figure 6.**
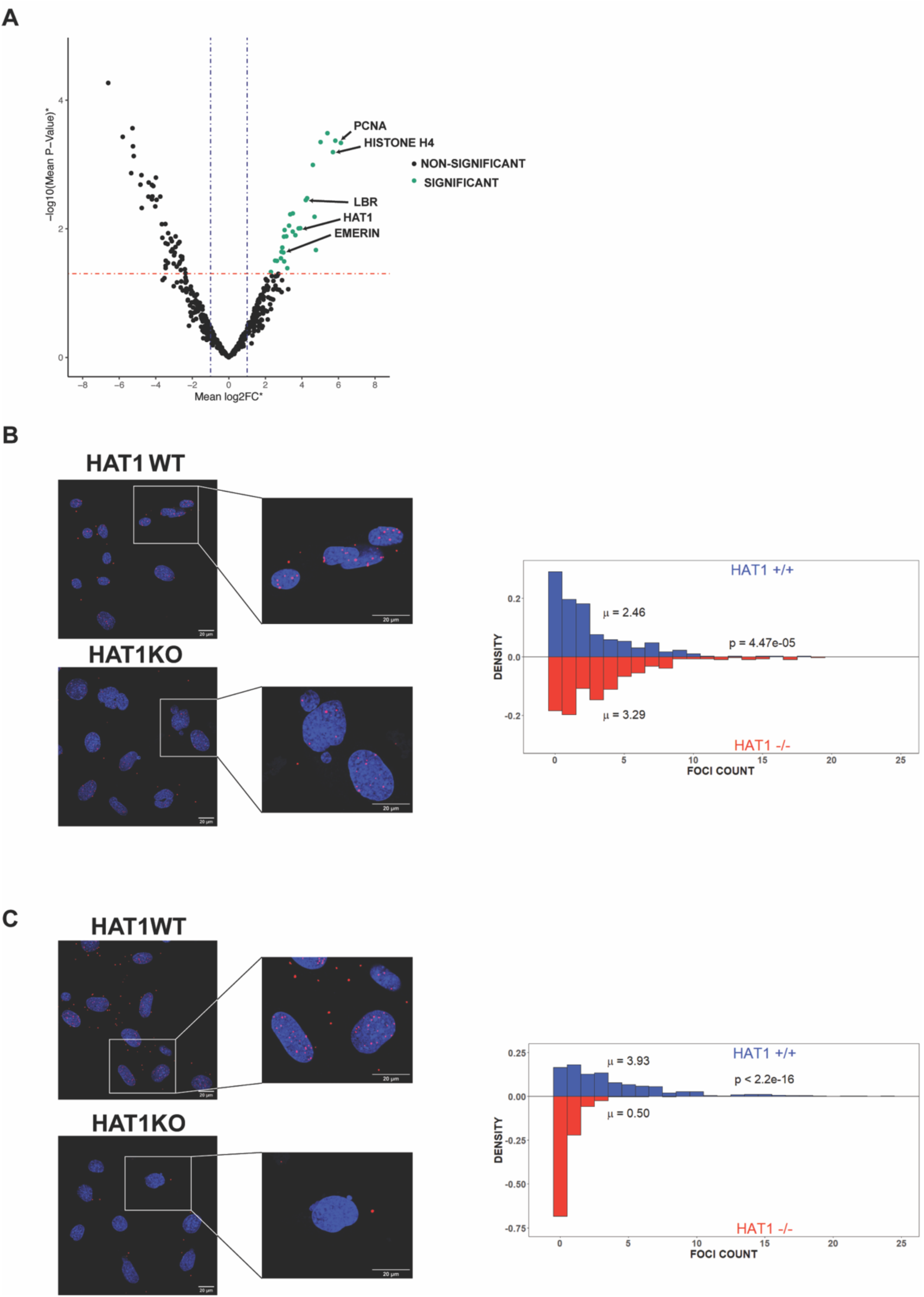
Hat1 colocalizes with nuclear lamina proteins. (A) Volcano plot showing average detected abundance and statistical significance of proteins identified by the Hat1-APEX2 construct after filtering with the control samples. (B) PLA between two Lamin B1 antibodies. Example images (left) and distribution of foci counts per cell (right) in Hat1 KO and WT cells. P-values calculated by Kolmogorov-Smirnov test. (C) Example images (left) and distribution of foci counts (right) of PLA between Hat1 and Lamin B1 in Hat1 KO and WT cells. P-values calculated by Kolmogorov-Smirnov test.

We next used a proximity ligation assay (PLA) to directly visualize the association of HAT1 with the nuclear lamina in cells. PLAs determine whether two molecules reside close to each other in the cell by employing two species-specific secondary antibodies that are fused to oligonucleotides. If the secondary antibodies recognize primary antibodies that are in close proximity, the oligonucleotides can both bind to a nicked circular DNA, creating a template for rolling circle replication. This amplifies sequences that can be bound by a fluorescent probe and visualized. As a positive control, we performed a PLA using two different antibodies that each recognize lamina B1. As seen in Figure 6B, a positive PLA signal is detected in both HAT1 WT and HAT1 KO iMEFs. We then combined one of the lamin B1 antibodies with an antibody recognizing HAT1. PLA signals are readily detected in HAT1 WT iMEFs but are lost in HAT1 KO iMEFs, confirming that HAT1 localized to the nuclear lamina (Figure 6C).

## Discussion

The reestablishment of heterochromatin after S-phase is a gradual process which takes the better part of a cell cycle (27,51,77). In recent years, studies have begun to reveal the mechanisms involved. Suv39h1/2 and HP1 mutually promote one-another’s retention at heterochromatin, leading to transcriptional silencing, deposition of H3K9me3, and the recruitment of chromatin remodelers, HDACs, histone methyltransferases, and more (78-83). Many factors associated with the replication fork are essential for heterochromatin regulation because of their role in recycling parental histones(84-86). Chromatin assembly factor 1 (CAF-1) also plays a crucial role by depositing new histones to prevent nucleosome depletion across the genome(87,88). However, an active role for new histone modifications in controlling chromatin re-establishment has not been recognized.

Using custom Hat1 KO cell lines which lack nascent H4K5/12ac, we show that this loss leads to complex changes in lamina-associated chromatin, with a net shift towards more repressive states. Lack of Hat1 causes an enrichment of Suv39h2 and corresponding depletion of G9a in LADs, which leads to increased H3K9me3 and loss of H3K9me2.

Interestingly, not all LADs respond to Hat1 loss in identical fashion. While almost all experience some reduction in histone acetylation, the shift to a stronger constitutive heterochromatin state is particularly seen in a subset of LADs, many of which were previously identified as HADs. While LADs in iMEFs can be defined by enrichment of either H3K9me3, H3K9me2, or H3K27me3, HADs fall almost entirely into the first of these categories. HADs are enriched in H3K9me3 and HP1β, depleted in H3K27me3, and above average size for LADs. It appears that the increase in Suv39h2 is taking place largely in regions where it was already enriched, and we are observing enhanced, not *de novo* recruitment. However, H3K9me3 abundance is not sufficient to explain why only this subset of LADs gains Suv39h2, as there are LADs with just as high or higher Suv39h2 levels which do not gain it upon Hat1 KO. We also see inconsistency in whether or not LADs gain HP1 upon Hat1 loss. We only examined the HP1β isoform. It would be interesting to see whether the other two HP1 isoforms increase in heterochromatin, particularly HP1α, which, like HP1β, is enriched in chromocenters (89). We were unable to determine a single structural feature which would explain the greater sensitivity of some LADs to Hat1 loss. Numerous factors, including the histone PTM signature, DNA methylation, 3D chromatin interactions, lamina-association, transcription, or sequence-specific DNA binding proteins, may distinguish them from other LADs, and it is likely that there are multiple contributing factors.

Suv39h1 and Su39h2 share the same general structure apart from an 81 amino acid N-terminal basic domain which is present only in Suv39h2 (90,91). As a result, they are often treated as interchangeable, though numerous studies have revealed differences in their substrate specificity, strength of localization to heterochromatin, and silencing capacity (92-94). One study found that Suv39h2 is primarily responsible for the deposition of H3K9me3 at constitutive heterochromatin and localizes to chromocenters more independently than Suv39h1 does, while Suv39h1 contributes more to transcriptional silencing(94). Our finding that only Suv39h2 is enriched in constitutive heterochromatin while Suv39h1 is more enriched in genic regions is mostly consistent with this. However, immunofluorescence has suggested both isoforms to be enriched in chromocenters, without significant localization to the nuclear periphery. This may point to differences in Suv39h1 and Suv39h2 localization between different cell types or cell lines. The unique N-terminal domain of Suv39h2 can bind RNA. Major satellite repeat transcripts help recruit it to pericentric regions but may also reduce its activity in some contexts (91,95). It would be interesting to test whether non-coding RNA transcripts contribute to Suv39h2 recruitment in our MEFs, though increased transcription upon loss of H4K5/12ac would be unexpected.

The observed loss of acetylation in LADs upon Hat1 KO is interesting since LADs are generally perceived as lacking histone acetylation. There are at least two ways to interpret this finding which are not mutually exclusive: (1) LADs maintain some basal level of histone acetylation throughout the cell cycle, possibly in order to preserve plasticity and the potential for activation, and thus are regulated by opposing pathways that preserve heterochromatin while ensuring that they are not as heterochromatic as they could be. (2) A specific subpopulation of cells possesses histone acetylation in LADs but cannot maintain it in the absence of Hat1. The second interpretation is consistent with S/early G2 phase cells being enriched in acetylated, newly synthesized histones. We have previously shown that nascent H4K5/12ac is eliminated in Hat1 KO cells and that Hat1 loss indirectly leads to depletion of H3K9 and K27 acetylation on nascent chromatin (58). While H4K5/12ac acetylation is probably reduced genome-wide in these cells, the relative effect is probably much more pronounced in heterochromatin, where parental histones are rarely acetylated, than in euchromatin, where acetylation is already enriched on many parental histones.

There are several ways in which acetylation of nascent chromatin might regulate heterochromatin maturation. H4K5/12ac can recruit bromodomain proteins such as ATAD2, which competes with HDACs 1 and 2 and influences HP1 recruitment, or BRPF3, which complexes with the H3K14 acetyltransferase KAT7 (96-99). We previously found that without Hat1, nascent chromatin was depleted in Brd3 and Brg1, both readers of acetylation on H4 K5 and/or K12 (61). It is also possible that H4K5/12ac affects the production of nascent chromatin transcripts, which helps recruit regulatory proteins and promote chromatin maturation(100-102). Hat1 or H4K5/12ac on new histones may even directly influence the recruitment or catalytic activity of H3K9 methyltransferases. As suggested above, Hat1 may also regulate the presence of other acetylation marks on nascent chromatin besides H4K5/12ac, which could have numerous downstream effects. Acetylation of H3K14 or H3K18 is found on over 20% of new histone H3 in human cells, and it is unknown whether they are deposited across the genome or biased toward specific regions(52). Ubiquitylation of H3K18 and H3K14, especially the latter, can promote H3K9me3 by Suv39h1/2, and their acetylation would block this ubiquitylation (46,47,103,104). In this context, it is interesting that the pattern and Hat1-sensitivity of H3K14ac is markedly different between HADs and nhLADs. H3K9ac could directly slow the spread of heterochromatin by reducing the amount of K9 available to be methylated. Since H3K9me3 must exceed a critical density in order to be propagated by the read-write mechanism, it is possible that small differences in H3K9ac could dramatically affect H3K9 spreading, especially near the threshold level (44).

Constitutive heterochromatin spreads via a positive feedback loop in which H3K9me3 recruits repressors such as HP1 proteins or Suv39h1/2 which promote the deposition of more H3K9me3 on nearby nucleosomes. The increased H3K9me3 observed in Hat1 KO HADs raises the question of why constitutive heterochromatin spreading into neighboring euchromatin is not observed. LAD boundaries remain remarkably consistent in the absence of Hat1, even those which gain H3K9me3 and Suv39h2. A potential explanation is the increased H4K5ac observed just outside HAD/nhLAD borders in Hat1 KO cells. This suggests that a protective mechanism exists in cells that senses the chromatin state of LAD borders and can respond to increased H3K9me3 by the recruitment of other HATs that prevent heterochromatin spreading.

It is uncertain what fraction of nucleosomes are trimethylated in a particular region of the genome. Our results suggest that nucleosomes in HADs are not saturated with H3K9me3, given the increase observed upon Hat1 KO. It is interesting to note, however, that LADs which gain H3K9me3 in Hat1 KO cells tend have much lower WT levels of H3K9me3 on average than those in which H3K9me3 is unchanged or lost (Figure S7). This suggests that LADs which are closest to saturating levels of H3K9me3 are less likely to gain more upon Hat1 loss. Alternatively, we may be observing increased H3K9me3 in a subset of cells which had lower levels than the rest of the population.

Despite the numerous chromatin changes caused by loss of Hat1, the changes in gene expression are relatively few (Popova et al 2021), probably because most of the chromatin changes take place in very gene poor regions. This seems to conflict with the vital importance of Hat1 for mammalian development. We hypothesize that Hat1 must moderate heterochromatin re-establishment to allow binding of lineage-specific transcription factors and facilitate proper differentiation during animal development. This is supported by our recent findings that Hat1 loss interferes with differentiation of intestinal stem cells *in vivo* and *in vitro,* and produces enhanced domains of H3K9me3 analogous to those described in this study (Nagarajan et al BIORXIV/2026/712164).

While many interesting questions remain, the results of this study supply an important advance in our understanding of Hat1’s role in epigenome regulation. Hat1 contributes to the regulation of H3K9 methyltransferase recruitment and regulates the balance of active and repressive features in heterochromatin, making it a vital regulator of chromatin inheritance.

## Data Availability

ATAC-seq data, HAD locations and LAD locations used for our analysis were previously published and are deposited in Gene Expression Omnibus GSE178592 and GSE281928 (64,65). All data generated in this study has been deposited in the NCBI Gene Expression Omnibus under accession numbers GSE325188 (CUT&Tag), GSE325189 (CUT&RUN), GSE325190 (ChIP-seq), and GSE325191 (RNA-seq).

## Funding

This work was supported by grant R01 GM144601 to MRP.

## Notes

### Competing Interest Statement

The authors have declared no competing interest.

