## Supplemental Material for "Hat1 Orchestrates Heterochromatin Inheritance by Regulating Localization of H3K9 Methyltransferases"

### Supplemental Figure Legends

**Figure S1**. H3K4me1 and H3K27ac in inter-LADs. (A) IGV browser comparison of H3K4me1 and H3K27ac over an ~8.5 Mb inter-LAD region on chromosome 1. (B-C) Heatmaps of H3K4me1 and H3K27ac across a 30 kb window centered around enhancers genome-wide.

**Figure S2**. IgG control in Hat1 KO and WT cells. (A) Boxplot of Hat1 KO/WT ratios for acetylation and IgG in all LADs (***: $p < 0.001$, **: $p < 0.01$, *: $p < 0.05$). (B) Same as A, specific for HADs.

**Figure S3**. HP1 western blots and quantitation of heterochromatin changes. (A) log2 KO/WT fold-changes of H3K9me3 (top left), H3K9me2 (top right), H2K27me3 (bottom left), and HP1β (bottom right) in LADs, colored by whether or not the LAD overlaps a HAD. (B-D) Western blots for (B) HP1α, (C) HP1β, and (D) HP1γ in separate iMEF cell lines derived from 3 WT and 3 Hat1 KO mice. (E) Regression of the log2 KO/WT fold-change of H3K9me3 vs. that of HP1β in HADs. Each LAD is a single data point.

**Figure S4**. HMT western blots and quantitation in LADs. (A-C) Western blots for (A) Suv39h1, (B) G9a, Suv39h2, and (C) Setdb1 in separate iMEF cell lines derived from 3 WT and 3 Hat1 KO mice. (D) Regression of G9a vs H3K9me2 levels in LADs (left) and inter-LADS (right). To allow comparison between LADs and iLADs, signal was summed in 10 kb bins and those with at least 50% LAD overlap were considered LAD bins while others were considered as iLAD bins. Due to the number of bins, the plot was divided into a fine grid and boxes colored by the number of points overlapping it. (E) Regression analysis of G9a vs Suv39h2 levels in LADs and iLADs, as in panel D. (F) log2 KO/WT fold-changes of Suv39h1 (top left), Suv39h2 (top right), Setdb1 (bottom left), and G9a (bottom right) in LADs, colored by whether or not the LAD overlaps a HAD.

**Figure S5**. Linear regression of HMTs and H3K9me2/3. (A-E) Regression of the $\log_2$ KO/WT fold-change for the indicted features across all LADs. Each LAD is a single data point

**Figure S6.** Western blot for HEK293T cells transfected with APEX2-HAT1, APEX2 only, and untransfected cells. Cytoplasmic extract (left) and nuclear extract (right) were probed separately following subcellular fractionation.

**Figure S7**. H3K9me3 profile grouped by H3K9me3 change. Profile and heatmap of H3K9me3 across all LADs in Hat1 WT cells, grouped by whether they lose, gain, or see no change in H3K9me3 in Hat1 KO cells (based on 1.25-fold change threshold). Profiles show all LADs scaled to the same size and extend 300 kb beyond each LAD border.

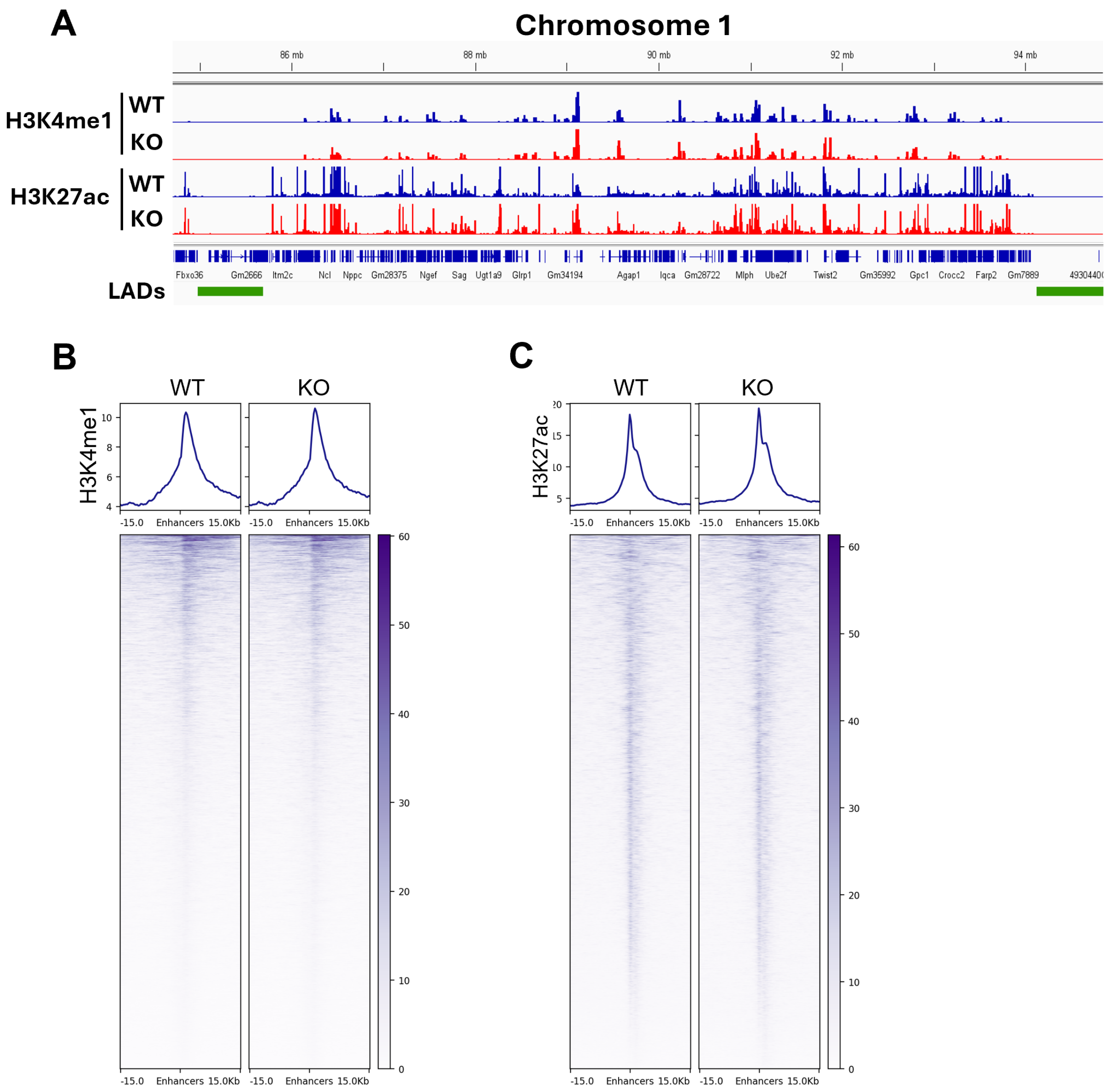
A
Chromosome 1
86 mb
88 mb
90 mb
92 mb
94 mb
H3K4me1
WT
KO
H3K27ac
WT
KO
Fbxo36 Gm2666 Itm2c Ncl Nppc Gm28375 Ngef Sag Ugt1a9 Glrp1 Gm34194 Agap1 Iqca Gm28722 Mlph Ube2f Twist2 Gm35992 Gpc1 Crocc2 Farp2 Gm7889
LADs
B
WT
KO
H3K4me1
-15.0 Enhancers 15.0Kb
C
WT
KO
H3K27ac
-15.0 Enhancers 15.0Kb

**Figure S1**

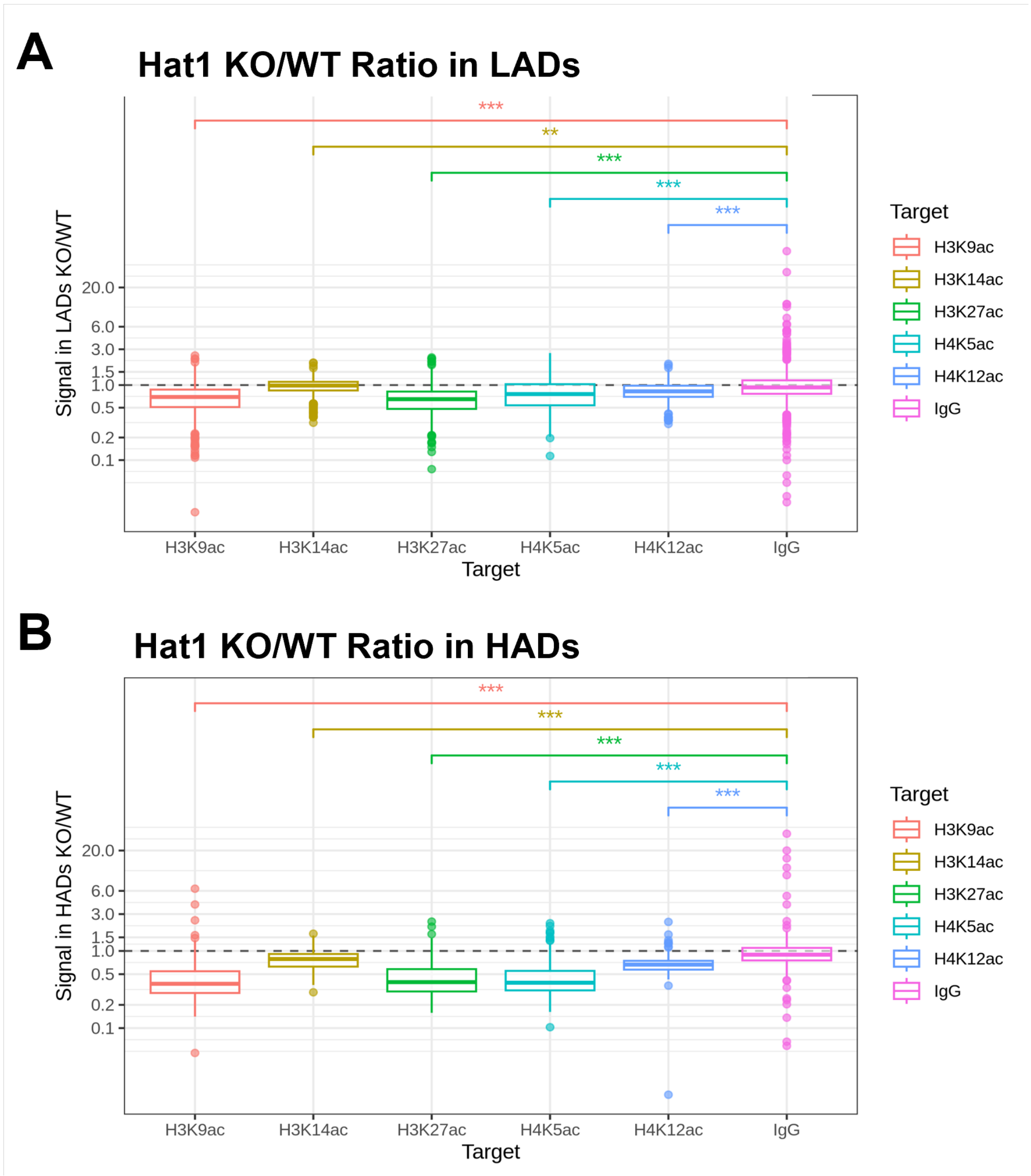
A
Hat1 KO/WT Ratio in LADs
***
**
***
***
***
Signal in LADs KO/WT
20.0
6.0
3.0
1.5
1.0
0.5
0.2
0.1
H3K9ac
H3K14ac
H3K27ac
H4K5ac
H4K12ac
IgG
Target
Target
H3K9ac
H3K14ac
H3K27ac
H4K5ac
H4K12ac
IgG
B
Hat1 KO/WT Ratio in HADs
***
***
***
***
***
Signal in HADs KO/WT
20.0
6.0
3.0
1.5
1.0
0.5
0.2
0.1
H3K9ac
H3K14ac
H3K27ac
H4K5ac
H4K12ac
IgG
Target
Target
H3K9ac
H3K14ac
H3K27ac
H4K5ac
H4K12ac
IgG

**Figure S2**

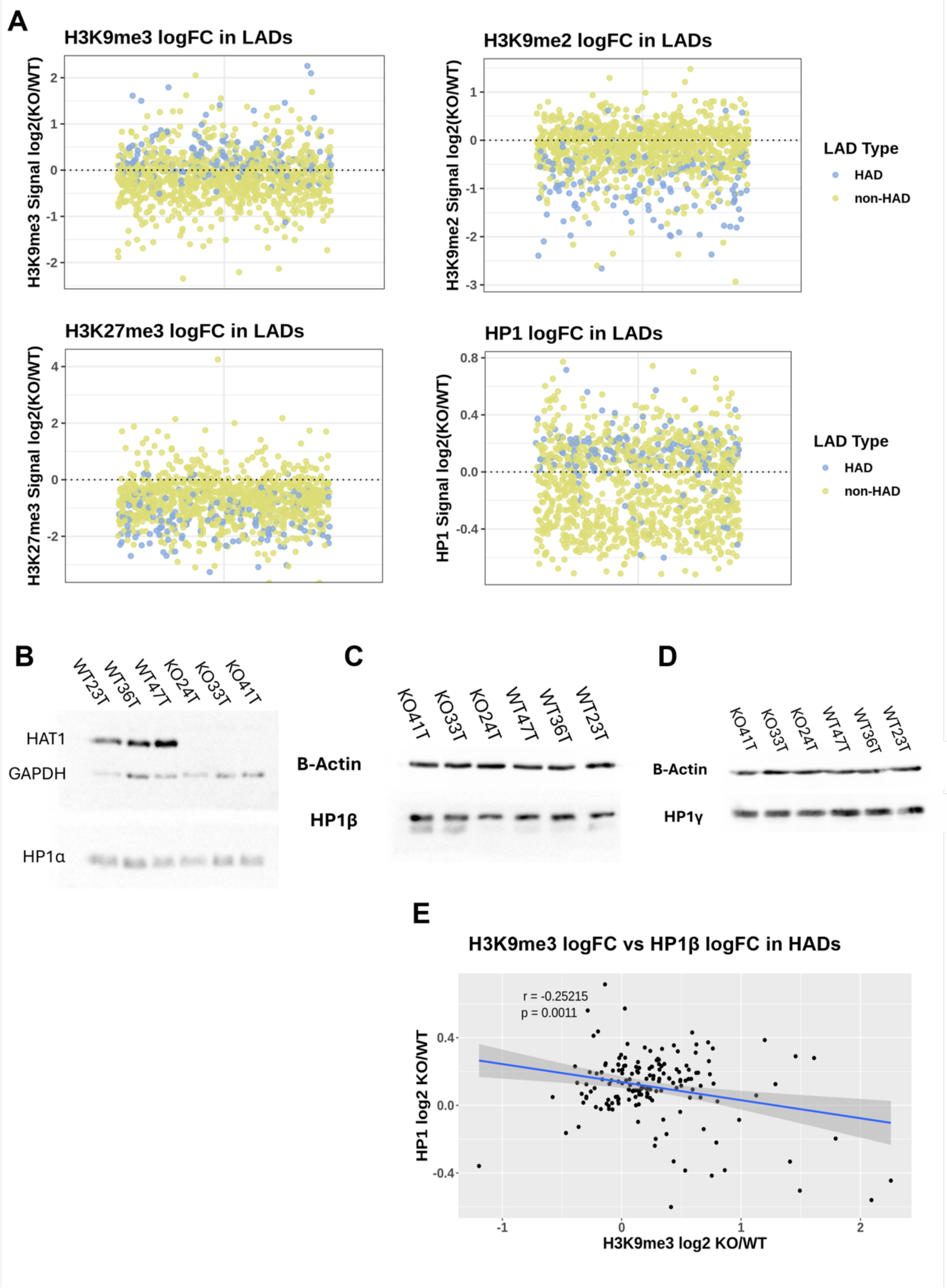
A
H3K9me3 logFC in LADs
H3K9me3 Signal log2(KO/WT)
H3K9me2 logFC in LADs
H3K9me2 Signal log2(KO/WT)
LAD Type
HAD
non-HAD
H3K27me3 logFC in LADs
H3K27me3 Signal log2(KO/WT)
HP1 logFC in LADs
HP1 Signal log2(KO/WT)
B
WT23T
WT36T
WT47T
KO24T
KO33T
KO41T
HAT1
GAPDH
HP1α
C
KO41T
KO33T
KO24T
WT47T
WT36T
WT23T
B-Actin
HP1β
D
HP1γ
E
H3K9me3 logFC vs HP1β logFC in HADs
r = -0.25215
p = 0.0011
HP1 log2 KO/WT
H3K9me3 log2 KO/WT

**Figure S3**

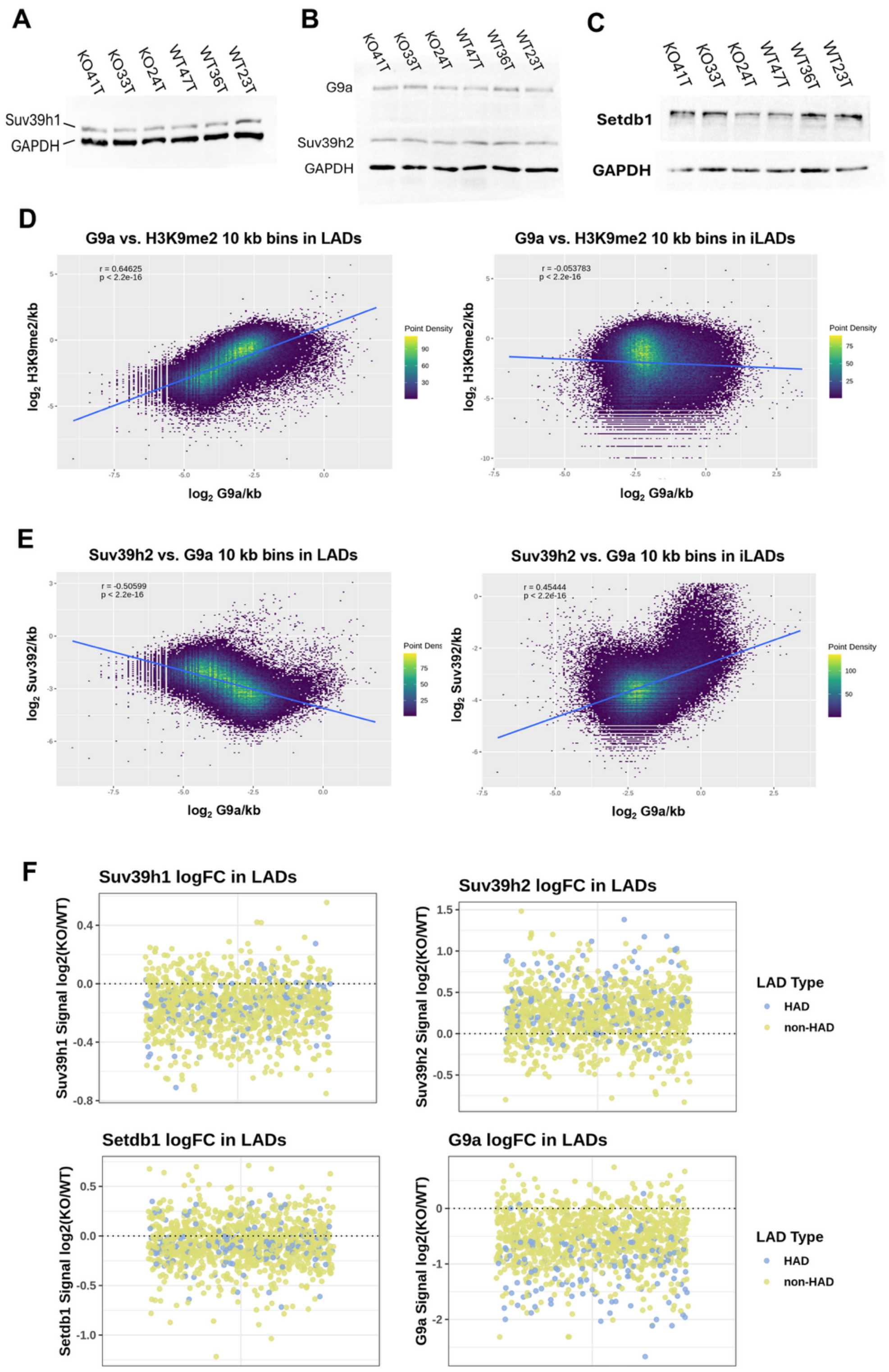
A
KO41T
KO33T
KO24T
WT47T
WT36T
WT23T
Suv39h1
GAPDH
B
G9a
Suv39h2
GAPDH
C
Setdb1
GAPDH
D
G9a vs. H3K9me2 10 kb bins in LADs
r = 0.64625
p < 2.2e-16
log2 H3K9me2/kb
log2 G9a/kb
Point Density
G9a vs. H3K9me2 10 kb bins in iLADs
r = -0.053783
p < 2.2e-16
E
Suv39h2 vs. G9a 10 kb bins in LADs
r = -0.50599
p < 2.2e-16
log2 Suv392/kb
Suv39h2 vs. G9a 10 kb bins in iLADs
r = 0.45444
p < 2.2e-16
F
Suv39h1 logFC in LADs
Suv39h1 Signal log2(KO/WT)
Suv39h2 logFC in LADs
Suv39h2 Signal log2(KO/WT)
LAD Type
HAD
non-HAD
Setdb1 logFC in LADs
Setdb1 Signal log2(KO/WT)
G9a logFC in LADs
G9a Signal log2(KO/WT)

**Figure S4**

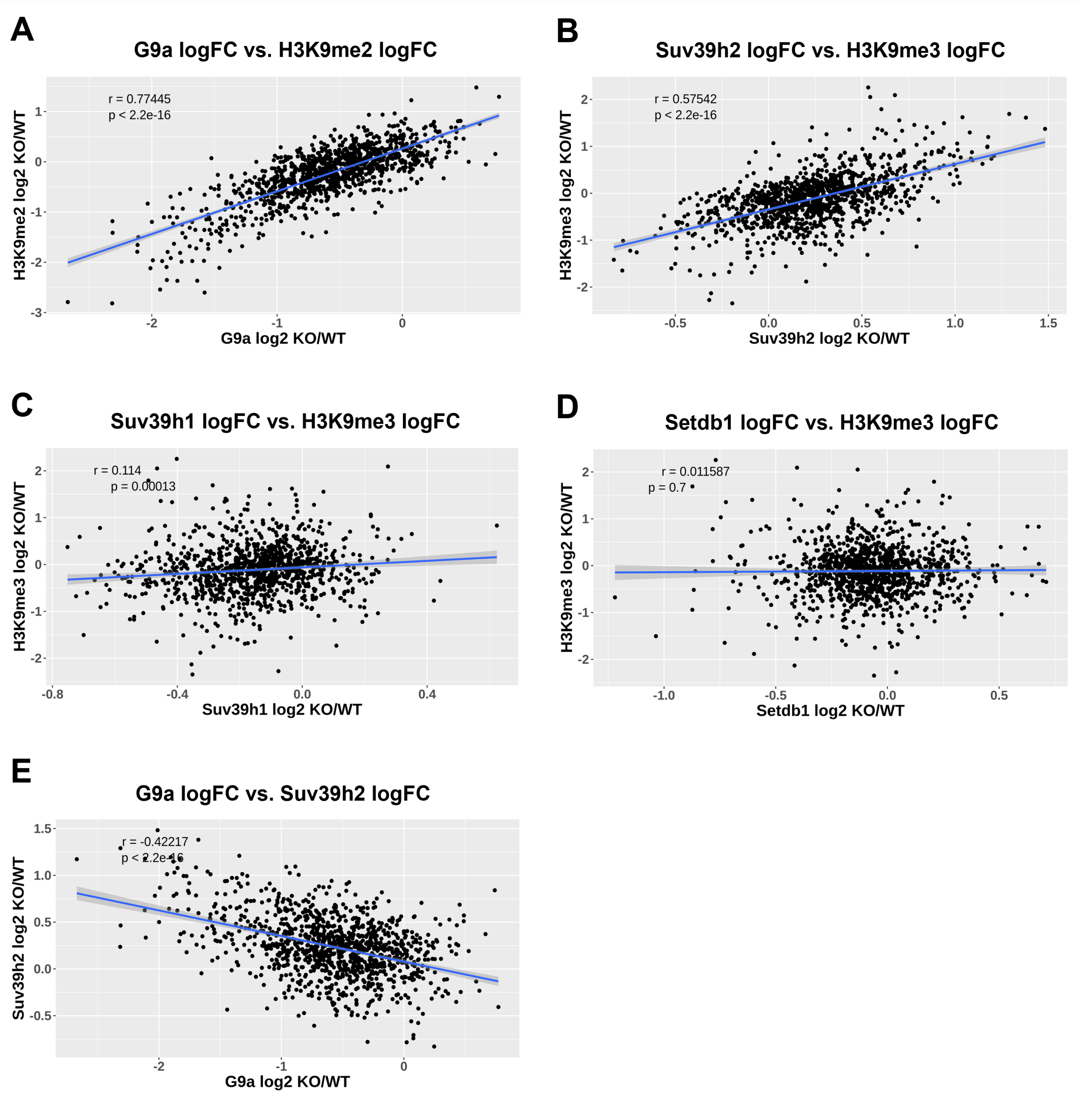
A
G9a logFC vs. H3K9me2 logFC
r = 0.77445
p < 2.2e-16
H3K9me2 log2 KO/WT
G9a log2 KO/WT
B
Suv39h2 logFC vs. H3K9me3 logFC
r = 0.57542
p < 2.2e-16
H3K9me3 log2 KO/WT
Suv39h2 log2 KO/WT
C
Suv39h1 logFC vs. H3K9me3 logFC
r = 0.114
p = 0.00013
H3K9me3 log2 KO/WT
Suv39h1 log2 KO/WT
D
Setdb1 logFC vs. H3K9me3 logFC
r = 0.011587
p = 0.7
H3K9me3 log2 KO/WT
Setdb1 log2 KO/WT
E
G9a logFC vs. Suv39h2 logFC
r = -0.42217
p < 2.2e-16
Suv39h2 log2 KO/WT
G9a log2 KO/WT

**Figure S5**

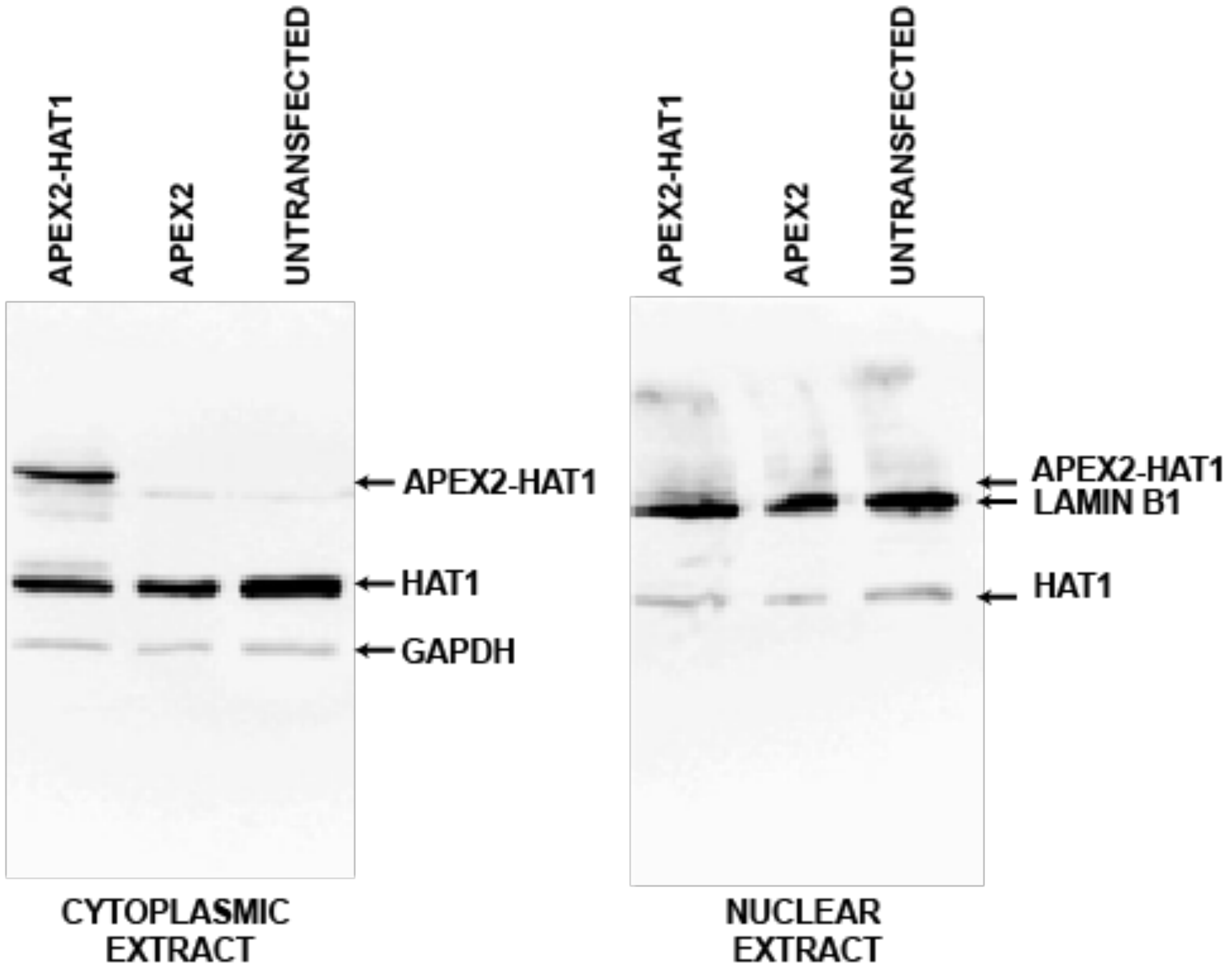
APEX2-HAT1
APEX2
UNTRANSFECTED
APEX2-HAT1
HAT1
GAPDH
CYTOPLASMIC EXTRACT
APEX2-HAT1
APEX2
UNTRANSFECTED
APEX2-HAT1
LAMIN B1
HAT1
NUCLEAR EXTRACT

**Figure S6**
